# Mechanisms of host exploitation by a microsporidian parasite

**DOI:** 10.1101/2024.12.13.628237

**Authors:** Luís M. Silva, Armelle Vallat, Jacob C. Koella

## Abstract

Parasites exploit their hosts to enhance their growth and reproduction, yet the mechanisms underlying host manipulation remain understudied for many taxa. The microsporidian *Vavraia culicis*, a potential biological control agent for mosquitoes, serves as an excellent model to explore such mechanisms. In this study, we investigate how infection by *V. culicis* lines that vary in virulence alters resource dynamics within the mosquito host *Anopheles gambiae*. Using metallomics and quantification of protein, carbohydrate, and lipid content, we show that infection alters host resource concentrations in ways that depend on parasite virulence. More virulent parasites led to increased protein levels and greater energy demands, evidenced by higher carbohydrate reserves. Additionally, infection with *V. culicis* impacted host metal content, particularly zinc and manganese, used by *V. culicis* independently of its evolutionary background. Iron availability, a key nutrient for parasite growth, enhanced spore production, with selected parasite lines better able to exploit host iron than unselected. These findings provide insight into the mechanisms by which *V. culicis* manipulates host resources, shedding light on the role of host exploitation in parasite virulence and the potential use of microsporidia as biological control agents in vector biology.

## 1. Introduction

The infection of hosts and their consequent exploitation by parasites is a ubiquitous feature of life [1]. The negative impact a parasite has on its host, particularly on host survival, is commonly referred to as virulence [2]. Virulence arises from the interplay between host and parasite traits [3]. Hosts can influence the cost of infection by limiting parasite load or mitigating infection-induced damage - mechanisms known as resistance and tolerance, respectively [4–6]. Parasites, in turn, may exploit host resources to fuel their own growth (host exploitation or proliferation), or invest in toxins and other virulence factors that cause additional harm independent of parasite load (per-parasite pathogenicity or benevolence)[3,7,8].

Since parasites typically sequester resources from their hosts, many hosts have evolved strategies to restrict resource acquisition by parasites through both metabolic and behavioural changes [9]. The mechanisms limiting exploitation have been heavily studied in recent years and have given rise to nutritional immunity [10–13], where the main aim is to enhance host survival by manipulating the parasite’s ability to exploit the host. The specific resources targeted by parasites vary depending on the stage of infection (time) and the host tissues involved (space). However, most parasites universally require a ubiquitous set of resources. These range from dietary carbohydrates to essential minerals, which are often hijacked by dedicated parasite transporters. Among these, iron is perhaps the most intensively studied [14–17]. Its critical role in microbial metabolism and structural function is underscored by the widespread evolution of siderophores - iron-chelating molecules used by microbes to extract iron from hosts [18–20]. Iron also acts as a developmental cue in some parasites, such as *Plasmodium* [21–24]. For instance, *Plasmodium falciparum* requires high iron concentrations for gametocyte fusion and successful development within the mosquito vector.

Despite increasing insight into parasite resource demands and sequestration strategies, many gaps remain, especially for obligate intracellular parasites such as microsporidia. Microsporidia are single-celled fungi that infect a wide range of hosts, from humans to dipteran insects [25–30]. Despite their compact genomes [31–33], they exhibit broad ecological diversity, colonising both aquatic and terrestrial environments [34]. Although studied for over 150 years [27], microsporidia have recently gained renewed attention due to their ability to suppress *Plasmodium* infection in *Anopheles* mosquitoes during co-infection [35–39]. Two species, *Vavraia culicis* and *Microsporidia MB*, have emerged as potential malaria control agents [35,40,41], supported by experimental [39,42] and field studies [37], respectively. However, the mechanisms by which these parasites exploit their hosts remain poorly understood. Considering *V. culicis* seems to elicit a limited host response, regardless of the vector host [43,44], here we turned focus to resource competition as a potential method of *Plasmodium*-inhibition. To date, only one study has partially characterised *V. culicis* exploitation of *Anopheles* larvae [45], and another has described limited effects on adult mosquito protein, carbohydrate and lipid content [46].

Understanding how microsporidia exploit their hosts is not only of interest for host-parasite interactions but also for vector biology, which entails the development of vector control strategies. In light of the long developmental time *Plasmodium* requires within its vector host, any host capable of transmitting *Plasmodium* will inherently favour parasites that also persist longer. Long-lasting parasites, such as some *V. culicis*, may thus evolve enhanced resource sequestration abilities, potentially outcompeting or excluding co-infecting parasites such as *Plasmodium spp*. Indeed, a recent study by Silva and Koella (2024) subjected *V. culicis* to artificial selection for either: i) early transmission and less time within the host, hereby referred to as *Early* treatment (i.e., parasites that killed the host under 7 days); or ii) late transmission and more time within the host, hereafter *Late* treatment (i.e., parasites that killed the host after 20 days) **(Fig. 1)** [44,47,48]. The authors found that lines selected for late transmission, and consequently persistence within the host, were more virulent, developed faster, and produced higher spore loads throughout their infective cycle (and the mosquito’s life) than those in *Early* or the unselected stock population (referred to here as *Stock* treatment). However, this came at a cost: *Late* parasites were less viable outside the host, highlighting a trade-off between adaptation within and among hosts, in the outside environment [48].

**Figure 1.**
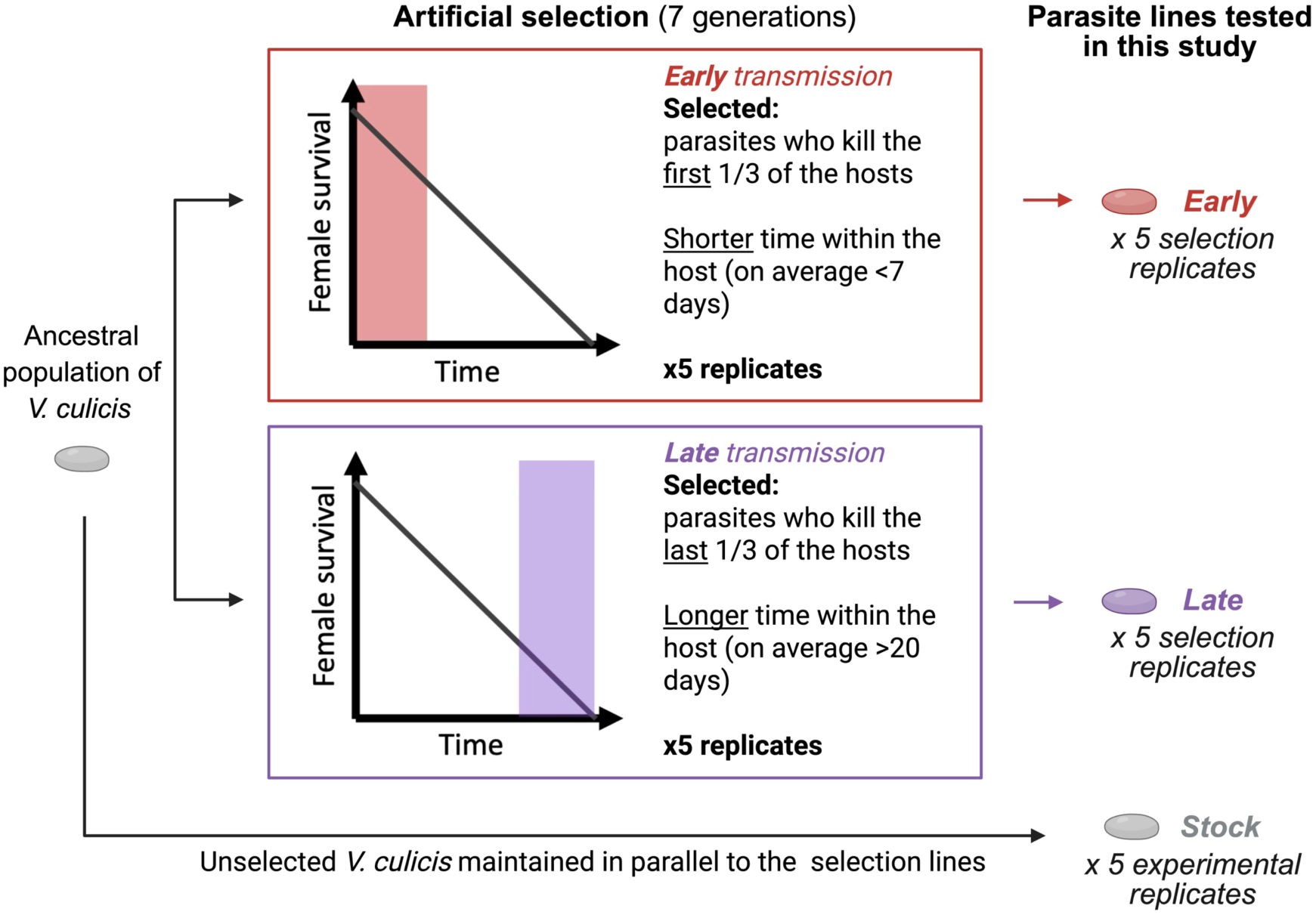
Illustration of the artificial selection experimental setup. An ancestral population of *V. culicis* was used to infect a Kisumu colony of the mosquito host *A. gambiae*. Two selection treatments were generated: i) *Early*, encompassing the parasites that were retrieved from the first third of the mosquitoes that died in the population; and ii) Late, including the parasites that were collected from the last third of the mosquitoes to die. The parasite went through seven serial passages before conducting the experiments in this study. Each selection treatment was replicated five times. As a baseline reference, the unselected parasite kept in a similar rearing parallel to the selection lines was also included in this study (i.e., *Stock*). To facilitate statistical comparisons, the *Stock* was also replicated five times. Further details can be found in Section 2.1 of the Materials and Methods and in the original study describing the selection lines, Silva and Koella (2024), bioRxiv.

In this study, we took advantage of these selected lines of *V. culicis* to: a) characterise *A. gambiae* exploitation by *V. culicis*; b) understand which mechanisms can be enhanced upon selection for early or late transmission (or less and more time within the host, respectively); and c) how changes in host exploitation relate to the different parasite lines’ virulence. To address these questions, we: i) quantified protein, carbohydrate, and lipid content; ii) performed metallomics; and iii) manipulated iron availability - known to influence *V. culicis* growth - by supplementing or depleting circulating iron in the adult mosquito. All measurements were taken on days 5, 10, and 15 of adulthood, following larval-stage infection with the selected (i.e., *Early* and *Late*) and unselected (i.e., *Stock*) *V. culicis* treatments **(Table 1)**. These time points correspond to three different phases of *V. culicis* growth: beginning, highest growth, and slowdown of the growth. We believe our findings will offer important insights into microsporidian biology, parasite evolution, and disease control, as the molecular machinery studied here is likely conserved or used by other parasites.

**Table 1.** Experiments conducted and respective replication. Experiments performed in this study (Figures 2 to 4), their treatments, and the number of replicates. The factor(s) of interest are the level at which conclusions are drawn. The scale at which the factor is applied represents the level at which treatments or factors are implemented, whereas the number of replicates at the appropriate scale shows the count of independent replicates for each treatment or factor level. Parasite treatment includes selected treatments, i.e., *Early* and *Late*, and the unselected *Stock*. Time consists of three time-points during mosquito adulthood, 5-, 10-, and 15-day post-adult emergence. Diet encompasses a standard sugar diet, supplemented with iron or chelated with BPS. The replication statement was adapted from the guidelines provided by the journal Functional Ecology (Wiley-Blackwell and British Ecological Society).

| Dependent variable(s) | Factor(s) of interest | Scale at which the factor is applied | Number of replicates at the appropriate scale |
| --- | --- | --- | --- |
| **Figure 2.** Protein, carbohydrate and lipid content throughout infection. |  |  |  |
| Protein, carbohydrate and lipid content | Parasite treatment and time | Individual mosquito | 10 mosquitoes per replicate (5) per time-point (3) |
| **Figure 3.** Metallomic profiles throughout infection. |  |  |  |
| Fe, Zn, Cu, Ca and Mn content | Parasite treatment and time | Pool of 10 mosquitoes | 1 pool per replicate (5) per time-point (3) |
| **Figure 4.** Effect of iron-supplementation and -chelation on parasite growth. |  |  |  |
| Spore load | Parasite treatment and diet | Individual mosquito | 10 mosquitoes per replicate (5) per time-point (3) per diet (3) |

## 2. Materials and Methods

### (a) Experimental model and mosquito rearing

We used the Kisumu strain of the mosquito host *Anopheles gambiae* (s.s.) and differently selected and unselected lines of the microsporidian parasite *Vavraia culicis floridensis* **(Fig. 1)**. To generate the selected lines, we used an ancestral population of *V. culicis* to infect naive *A. gambiae* mosquitoes. From the initial generation (F0), parasites were collected from the first one-third of mosquitoes to die (forming the *Early* treatment) and from the final one-third to die (forming the *Late* treatment). From F1 onwards, these parasite lines were maintained under the same selection protocol - *Early* and *Late* - but evolving independently. This selection process was repeated across seven generations for each of five replicate lines per parasite treatment, allowing only the parasite to evolve, not the host [47]. After seven generations, the selected parasite lines and an unselected reference line (*Stock*), maintained in parallel to the artificial selection experiment, were used in the described experiments. Further details on the selection experiment and its maintenance are described in Silva and Koella (2024). Experiments and rearing were conducted at standard laboratory conditions, 26 ± 1°C, 70 ± 5% relative humidity, and 12 h light/dark.

### (b) Experimental setup

All experiments were performed similarly **(Table 1)**. In each experiment, freshly hatched *A. gambiae* larvae were grown in groups of 50 per Petri dish (120 mm diameter x 17 mm) containing 80 ml of distilled water. Five petri dishes were prepared for each parasite treatment and replicate. Larvae in these dishes were fed daily with Tetramin Baby fish food according to their age: 0.04, 0.06, 0.08, 0.16, 0.32 and 0.6 mg/larva for 0, 1, 2, 3, 4 and 5 day-old or older larvae, respectively. Two-day-old larvae were exposed to 10,000 spores per larva of their respective *V. culicis* treatment. Pupae were then transferred to cages (21 x 21 x 21 cm) and left to emerge as adults. Each cage included individuals from all petri dishes within a given parasite treatment and replicate. In each experiment, alive mosquitoes were retrieved for the different assays. The collection was random within the same parasite treatment and replicate for all traits measured. For the entirety of their adult life, the mosquitoes were provided with a 6% sucrose solution up until trait measurement. Males were removed as soon as they emerged to avoid mating-associated costs.

### (c) Proteins, carbohydrates and lipids quantification

Mosquitoes were collected 5, 10, or 15 days after emergence and frozen at -80 °C. We quantified their proteins, carbohydrates, and lipid content through colourimetry, which we adapted from [49]. Protein, carbohydrates, and lipid concentrations were estimated from the same sample for each of the parasite treatments and respective replicates. Each individual body was weighed and then submerged in a 100 µL PBS pH 7.4 solution before homogenisation. To each sample, we added two stainless beads (Ø 2 mm) and homogenized the sample using a Qiagen TissueLyser LT at a frequency of 30 Hz for three minutes. Then, the protein, carbohydrate, and lipid content were assessed for each mosquito as described below. It is noteworthy that although we aimed to extract as much of the resource content from the mosquito, some still hold onto the fibrous tissue. Hence, our measurements concern content that is rather available and circulating, rather than structural.

Proteins were quantified by centrifuging the samples at 1100 RPM at 4° C for four minutes, adding 200 µL Bradford reagent to each solution, incubating at room temperature with soft agitation for 20 min, and reading the absorbance at 595 nm with a SpectraMax i3x plate reader. To quantify protein content, a standard curve of seven concentrations ranging from 400 µg/ml to 10 µg/ml of albumin in extraction buffer (100 mM monopotassium phosphate, 1 mM DTT, and 1 mM EDTA in Milli-Q water) was also prepared.

Carbohydrates were measured by first adding 20 µL of sodium sulphate at 20% to the homogenate, adding 5 µL of the extraction buffer and 1500 µL Chloroform-Methanol (1:2, v:v), vortexing, and centrifuging at 1100 RPM, 4°C for 15 minutes. The supernatant was then removed, of which 300 µL was transferred to a 96-well borosilicate plate and left to evaporate under the hood for approximately 30 minutes at room temperature. 240 µL of anthrone reagent was added to each sample and incubated at room temperature for 15 minutes. The plate was sealed and incubated in a water bath at 90°C for 15 minutes before measuring absorbance at 625 nm. A standard curve of seven concentrations ranging from 40 µg/ml to 1 µg/ml of glucose in Chloroform-Methanol (1:2, v:v) was also prepared to quantify carbohydrate content.

Lipids were measured by adding 200 µL of the supernatant to a 96-well borosilicate plate, then heated to 90 °C for 30 minutes. 10 µL of 98% sulfuric acid was added, and the sample was sealed and incubated for two minutes in a water bath at 90 °C. The samples were transferred to ice for three minutes before adding 190 µL of vanillin reagent. Samples were sealed and incubated at room temperature for 15 minutes with gentle agitation. Their absorbance was measured at 525 nm. A standard curve of seven concentrations ranging from 200 µg/ml to 5 µg/ml of triolein in Chloroform-Methanol (1:2, v:v) was also prepared.

A blank sample was also included in all the standard curves to control for contamination. To acquire individual resource content, the amount of resources measured was divided by the weight of the mosquito. Details regarding the experiment scale and replication can be found in **Table 1**.

### (d) Metals quantification

Five, 10 and 15 days after emergence, 30 alive females per parasite line and replicate were snap-frozen in a 2 ml microcentrifuge tube at -80°C. The samples were freeze-dried and submitted to a mineralisation procedure using a Titan MPS microwave (Perkin Elmer, Waltham, MA). Each sample was placed into a PTFE digestion vessel, and 2.5 mL HNO3 (Rotipuran Supra 69%, ROTH, Germany) and 2.5 mL H2O2 (analytical reagent grade, Fisher Scientific, UK) were added. The vessels were heated following a two-step digestion program consisting of a first step of 25 min until reaching 175°C and a second step of 10 min at 175°C. After cooling, the digested sample solutions were transferred to 15 mL Falcon PP tubes and stored in the fridge. A blank solution was prepared using the same procedure without the sample. The elements iron, zinc, copper, calcium and manganese were simultaneously analysed using inductively coupled plasma optical emission spectrometry (ICP-OES) [24]. An Avio 550 max equipped with a dual view and an S23 autosampler (PerkinElmer, Waltham, MA) was used. The instrument was operated in axial view with the following conditions: RF power 1450 W, cyclonic spray chamber with nebuliser nitrogen gas flow rate 0.7 mL/min, plasma argon gas flow rate 8 mL/min and auxiliary compressed air gas flow rate 0.2 mL/min. The sample was injected into the plasma at 1.30 mL/min. All the calibration curves were prepared in 2% HNO3 MilliQ water (w/w) in the concentration range of 0.1-10 mg/L and using SCP Science multi-element standards (SCP Science, France). Yttrium (Atomic spectroscopy standard, PerkinElmer Pure, 2% HNO3, 1000 ppm) was used as the internal standard and was added at a concentration of 1 mg/L in the blank standards and samples. The elemental wavelengths were 238.204 nm for Fe, 206.200 nm for Zn, 327.393 nm for Cu, 317.933 nm for Ca, 257.610 nm for Mn and 371.029 nm for Y. Measurements were performed in triplicates. Details regarding the experiment scale and replication can be found in **Table 1**.

### (e) Iron metabolism assay

In a separate experiment, after rearing larvae and pupae infected with their respective parasite treatment, all the pupae of a given combinatorial treatment (parasite treatment x replicate) were allowed to emerge within a cage. The emerging adults were then randomly split into three diet treatments, starting the experiment. Mosquitoes were fed with daily prepared cotton pads with either: i) iron (1 mM FeCl_3_ in 6% sucrose solution), ii) chelator (200 μM Bathophenanthroline disulfonic acid disodium salt (BPS) in 6% sucrose solution) or iii) or sugar (6% sucrose solution) for five days after emergence. From day six onwards, the mosquitoes were provided with cotton pads with only 6% sucrose solution. Alive individual mosquitoes were collected for spore load quantification 5, 10, or 15 days after emergence. They were placed individually into 2 ml microcentrifuge tubes with 1 ml of distilled water and a stainless-steel bead (Ø 5 mm, and homogenised with a Qiagen TissueLyser LT at a frequency of 30 Hz for one minute. The number of spores was counted in a haemocytometer under a phase-contrast microscope (400x magnification). Details regarding the experiment scale and replication can be found in **Table 1**.

### (f) Statistical analysis

All the analyses were performed with R version 4.3.1. Statistical analysis and their diagnosis were conducted using: “DHARMa” [50], “car” [51], “lme4” [52], “muhaz” [53], “emmeans” [54] and “multcomp” [55].

First, we tested if the parasite treatment or the day of collection affected the mosquito’s body weight. For this, we used a linear mixed model with log-transformed weight as a dependent variable and parasite treatment, time of collection and their interaction as explanatory variables and replicate as a random factor. Body weight was not affected by any of the factors (Parasite treatment: χ^2^ = 1.38, *df* = 2, *p* = 501; Time: χ^2^ = 0.80, *df* = 1, *p* = 0.373; Parasite treatment * Time: χ^2^ = 1.04, *df* = 2, *p* = 0.60).

Differences in resource content were assessed using a linear mixed model for each of the three resources (proteins, carbohydrates, and lipids), with parasite treatment and time of collection as explanatory variables and their interaction. Replicate was considered a random factor. Individual resource content was quantified by measuring the sample’s quantity and adjusting it according to the respective individual weight. The square root of each resource content was calculated to fit the model best.

We ran a linear mixed model for each metal element to test for differences in the metallomes of differently infected or uninfected mosquitoes across time. The dependent variable was the log-transformed concentration of the element obtained by ICP-OES, while the parasite treatment and assay day were explanatory variables and their interaction. Replicate was included as a random factor.

We assessed the effect of the parasite treatment and diet on spore rate and growth for each time point assayed. For each spore rate comparison, we ran a generalised linear mixed model with a binomial error structure, where parasite treatment and diet treatment were set as explanatory variables, their interaction, and the presence or absence of spores as a response variable. Replicate was included as a random factor. Concerning spore growth analyses, we only considered individuals with at least one spore. We then ran a linear mixed model with the same explanatory and random factors as the previous one.

## 3. Results

### (a) Resource and energy use in mosquitoes infected with lines of V. culicis

The protein content was affected by parasite treatment (χ^2^ = 9.75, *df* = 2, *p* = 0.008), time (χ^2^ = 45.28, *df* = 1, *p* < 0.001) and their interaction (χ^2^ = 24.75, *df* = 2, *p* < 0.001; Fig. 2a). In particular, protein content remained stable until day 10 of adulthood when it significantly increased for mosquitoes infected with late-selected spores (i.e., the most virulent ones) compared to the other parasite treatments (χ^2^ = 9.75, *df* = 2, *p* = 0.008).

**Figure 2.**
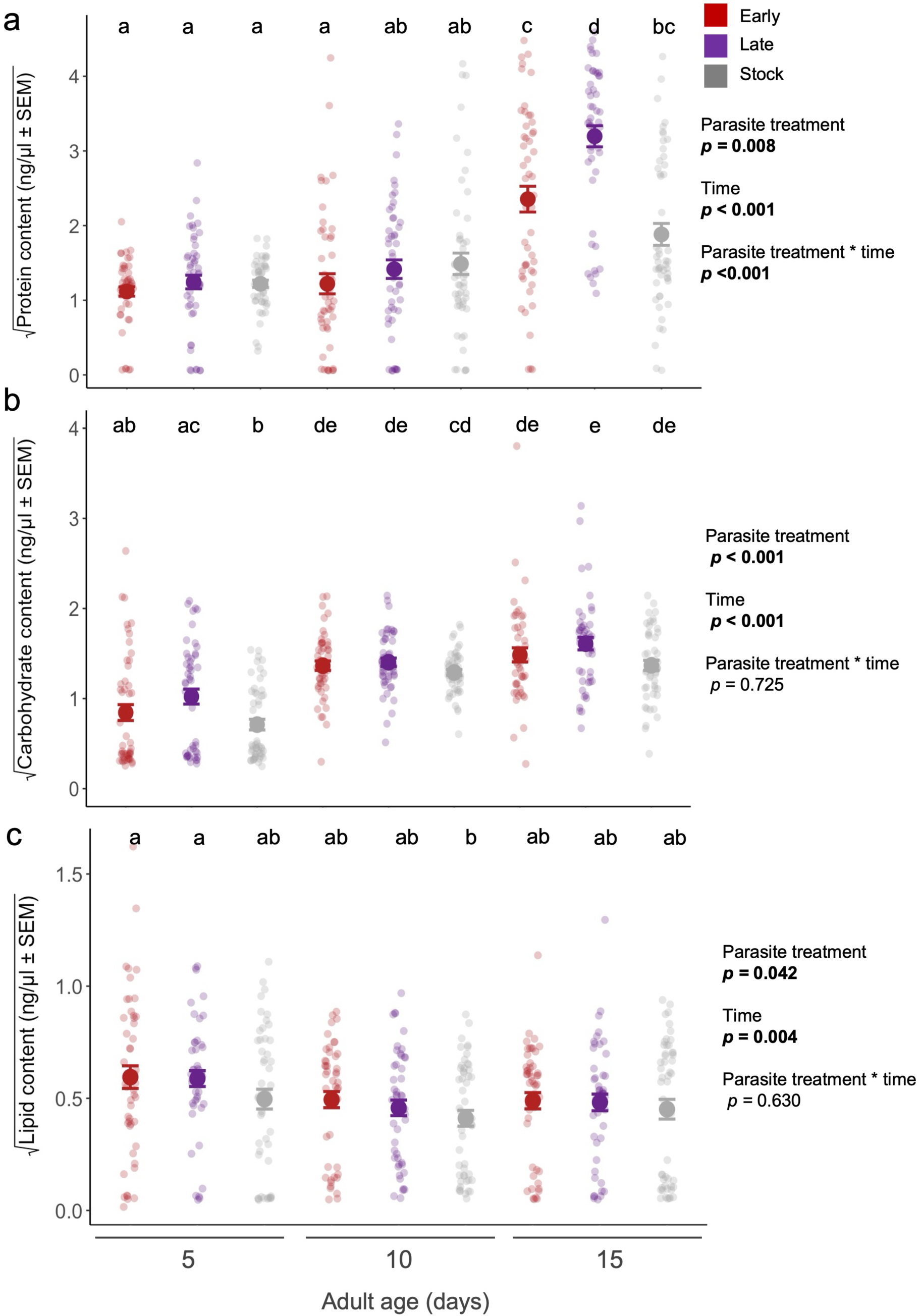
Mean protein, carbohydrate and lipid content through infection. Mean square root of the **(a)** protein, **(b)** carbohydrate and **(c)** lipid content at day 5, 10 and 15 of adulthood upon infection with early-, late-selected, or unselected stock parasite. Each time-point consists of individual resource content from 50 live females per parasite treatment. Statistical factors and respective *p-*values are shown on the right of each plot. Letters denote differences from multiple comparisons tests for an interaction between “parasite treatment” and “time”. For statistical details, see SI Appendix Table S1.

The carbohydrate content increased with infection time (χ^2^ = 139.65, *df* = 1, *p* < 0.001; Fig. 2b). It was higher in mosquitoes infected by the most virulent parasites (i.e., late-selected ones) than in those infected by the unselected or the early-selected ones (χ^2^ = 16.56, *df* = 2, *p* < 0.001).

The lipid content decreased with infection time (χ^2^ = 8.25, *df* = 1, *p* = 0.004) irrespective of treatment, and the mosquitoes infected with selected spores (i.e., early or late) had higher lipid content than the ones infected with the unselected parasites (χ^2^ = 6.33, *df* = 2, *p* = 0.042; Fig. 2c).

### (b) Metallomics of mosquitoes infected with lines of V. culicis

All metals (iron, copper, zinc, manganese) and calcium concentrations in the mosquito increased with time (Fe: χ^2^ = 18.88, *df* = 2, *p* < 0.001; Cu: χ^2^ = 11.93, *df* = 2, *p* = 0.003; Zn: χ^2^ = 18.62, *df* = 2, *p* = < 0.001; Mn: (χ^2^ = 17.91, *df* = 2, *p* < 0.001; Ca: χ^2^ = 34.08, *df* = 2, *p* < 0.001). While iron, copper, and calcium concentrations were similar for parasite-exposed and unexposed mosquitoes and for all parasite lines (Fe: χ^2^ = 2.26, *df* = 3, *p* = 0.521; Cu: χ^2^ = 0.66, *df* = 3, *p* = 0.88; Ca: χ^2^ =3.19, *df* = 3, *p* = 0.36), the zinc concentrations were lower in infected mosquitoes, irrespective of parasite line (Zn: χ^2^ = 13.97, *df* = 3, *p* = 0.003) and manganese concentration was higher (χ^2^ = 32.30, *df* = 3, *p* < 0.001) (**Fig. 3b, e**).

**Figure 3.**
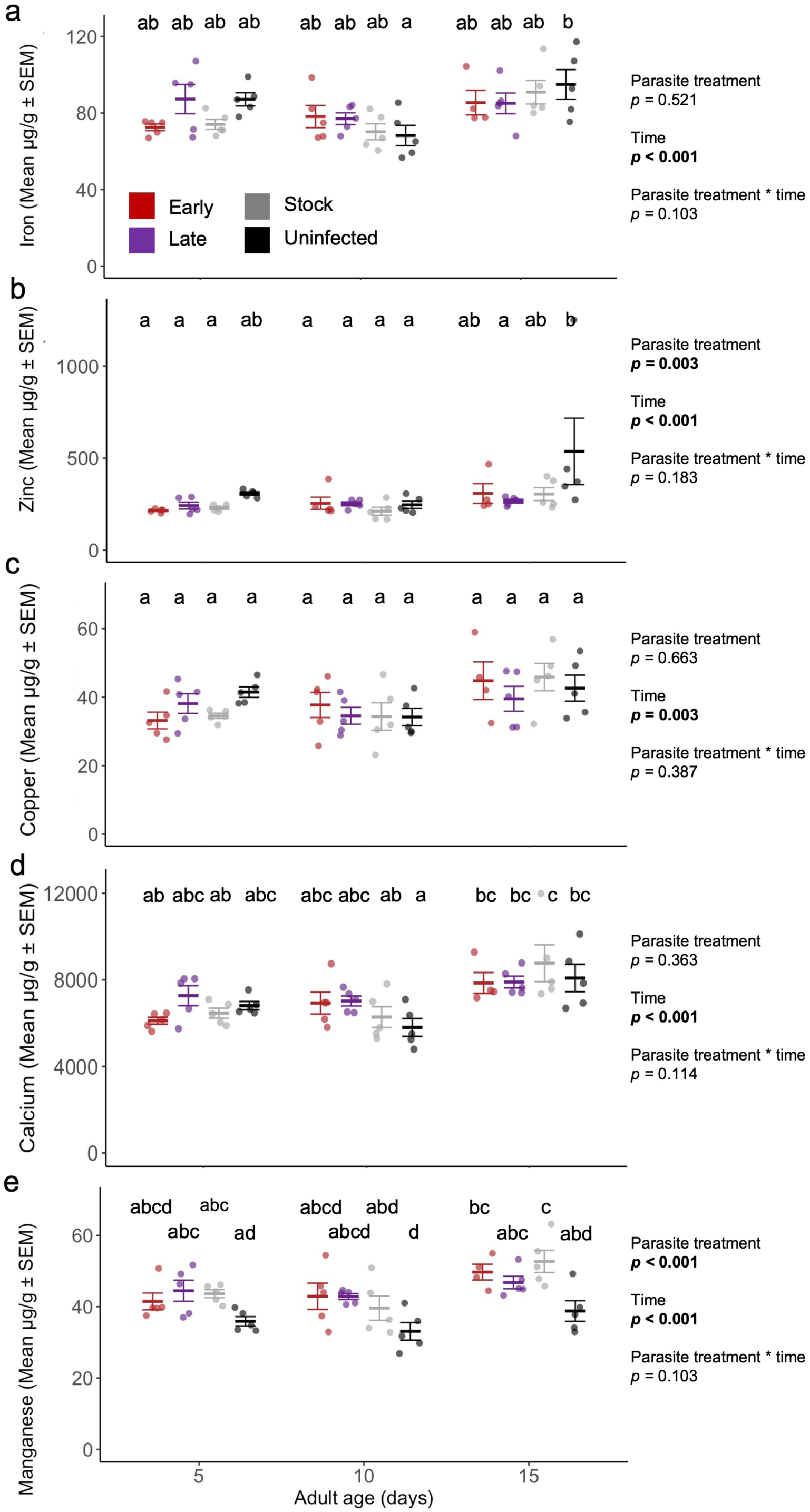
Metallomic profiles at adulthood. Metal concentrations determined by ICP-OES in uninfected *An. gambiae* and infected with one of three *V. culicis* parasite lines: early, late or unselected stock. Iron **(a)**, zinc **(b)**, copper **(c)**, calcium **(d),** and manganese **(e)** concentrations were quantified across adult age at days 5, 10 and 15 post emergences. For each time point, a pool of 30 females was used to assess metal concentration for each selection treatment x replicate combination (i.e., *n* = 5 data points per selection treatment). Statistical factors and respective *p-*values are shown on the right of each plot. Letters denote differences from multiple comparisons tests for an interaction between “parasite treatment” and “time”. The mean and standard error of the mean were plotted. For further statistical details, see SI Appendix Table S2.

### (c) Iron effect on parasite growth

In the iron supplementation or chelation experiment, all mosquitoes harboured spores by day 15. At days 5 and 10, the proportion of mosquitoes with spores was higher if the parasite had been late-selected than if it had been early-selected or not selected (day 5: χ^2^ = 15.43, *df* = 2, *p* < 0.001; day 10: χ^2^ = 8.46, *df* = 2, *p* = 0.015). On both days, iron supplementation increased while chelation decreased the spore rate (day 5: χ^2^ = 15.43, *df* = 2, *p* < 0.001; day 10: χ^2^ = 8.46, *df* = 2, *p* = 0.015). A similar pattern was observed for the spore load in individuals that harboured spores, with higher spore load after an iron-supplemented diet and lower spore load after a chelator-supplemented diet (day 5: χ^2^ = 12.45, *df* = 2, *p* = 0.002; day 10: χ^2^ = 37.60, *df* = 2, *p* < 0.001; day 15: χ^2^ = 79.65, *df* = 2, *p* < 0.001; **Fig. 4b**). Individuals infected with late-transmitted spores also showed the highest number of spores regardless of the time point or diet (day 5: χ^2^ = 37.75, *df* = 2, *p* < 0.001; day 10: χ^2^ = 37.60, *df* = 2, *p* < 0.001; day 15: χ^2^ = 26.73, *df* = 2, *p* < 0.001; **Fig. 4b**), except for time-point 15 days where infection by the unselected parasite reached the same load as the late-transmitted spores for the iron-supplemented diet (interaction between parasite and diet treatments: χ^2^ = 18.60, *df* = 4, *p* < 0.001; **Fig. 4b**).

**Figure 4.**
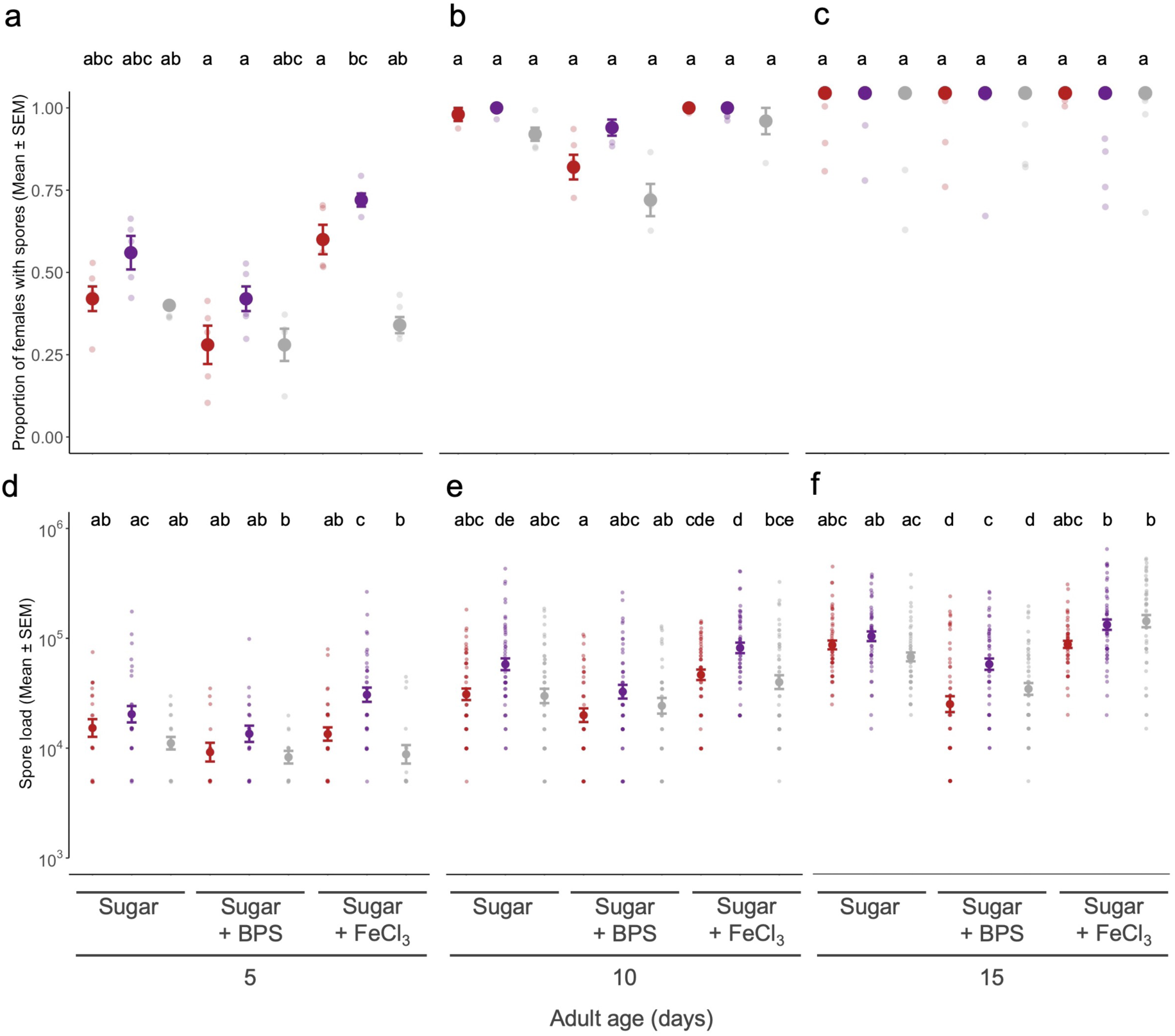
The effect of iron-supplementation and -chelation on parasite growth. The impact of a standard sugar diet (6% Sucrose), supplemented with an iron-chelator (6% Sucrose + 200 μl BPS) or iron itself (6% Sucrose + FeCl_3_ 1mM) on the presence of detectable spores (i.e., the proportion of females with spores secreted) **(a-c)** and spore load **(d-f)** of differently selected, or unselected, *V. culicis* treatments. Each selection treatment has a sample size of 50 females split by five experimental replicates for each time point. Letters denote differences from multiple comparisons tests for an interaction between “parasite treatment” and “diet” for each independent time point. For further statistical details, see SI Appendix Table S3.

## 4. Discussion

Parasites are masters at host exploitation, sequestering, and manipulating host resources to their benefit. In this study, we show how infection by the microsporidian parasite *V. culicis* alters host resource concentrations, particularly metals and energy sources stored in the mosquito host *A. gambiae*. We could also link virulence better to host exploitation by using parasite lines that differ in virulence - that is, the degree of harm inflicted on the host. The findings discussed below reveal microsporidia-specific mechanisms of host exploitation that may be inherently present or selectively enhanced under particular conditions. In addition to characterizing these mechanisms, we consider and discuss how their differential selection influences the potential role of microsporidia as biological control agents within vector biology.

### (a) Parasite virulence drives host energy use during infection

First, we observed that protein content in mosquitoes increased over time, particularly between days 10 and 15 of adulthood **(Fig. 2a)**. As most immune effectors and signalling pathways are protein-based, this increase may indicate a host shift to parasite recognition and consequent immune response. If so, our findings support the threshold hypothesis [56], which proposes that an immune response is triggered only once parasite load surpasses a certain threshold. This would explain the delayed rise in proteins and potential immune effectors, such as antimicrobial peptides.

Moreover, the magnitude of the increase varied with parasite virulence: infection with the most virulent parasite line (i.e., *Late*) led to higher host protein levels by day 15 **(Fig. 2a)**. A similar pattern has been observed in several other studies [7,57,58], and is likely linked to the much higher parasite burdens reached by *Late* parasites, as shown in our previous work [47]. These results also highlight the mosquito host’s ability to detect and adjust life history investments in immunity or other fitness traits, such as fecundity, based on the virulence of the infecting parasite. For instance, *Late* spores have been shown to induce earlier reproduction in *A. gambiae* mosquitoes [47].

We also measured two fundamental energetic reserves in mosquitoes: carbohydrates and lipids. Carbohydrates are typically more accessible and readily used, whereas lipids are a last resort or emergency energy source. Over time, we found that lipid reserves declined slightly **(Fig. 2c)**, while carbohydrate reserves increased for most parasite lines **(Fig. 2b)**. This pattern likely reflects our experimental conditions, where adult mosquitoes had ad libitum access to sucrose (a carbohydrate) but no lipid-based diet. Consequently, although both energy reserves are utilised during infection, as shown previously for this microsporidian species [46], carbohydrate stores are replenished, whereas lipid reserves are steadily depleted or sequestered. Interestingly, we do find a difference in energetic use from a previous study that only focused on infection by the unselected line of *V. culicis* [46]. While in this latter study, Zeferino and colleagues (2024) observed a shift in energy use later in mosquito life from carbohydrates to lipids, here we observed an almost immediate use of lipids and a gradual increase in carbohydrates, particularly in the selected parasite lines, *Late* and *Early*. Although perhaps speculative, we can hypothesize that this difference might be driven by the selection process, which selects for better-adapted parasites to *A. gambiae* and therefore, better host exploitation. To aggravate this, parasite lines that better exploit the host, such as *Early* and particularly *Late*, and produce higher numbers of infective spores, might have higher energy demands. Previous studies on this model system have shown the ability of this parasite to change the host behaviour [59,60]. Hence, it might be conceivable that either the host or the parasite is driving the mosquito to feed more and have its carbohydrates replenished for the duration of the infection.

Therefore, parasite treatment influenced the dynamics of these energy sources **(Fig. 2bc)**. Carbohydrates increased proportionally to parasite virulence, which can reflect the host’s increased energetic demand for immunity [61–63] and damage control [64], or the parasite’s needs, as mentioned above. Additionally, *V. culicis* draws upon the host’s carbohydrate reserves for its own consumption early in adulthood [46]. Thus, more virulent parasites, which elicit stronger immune responses and produce more infective spores, impose greater energy demands on the host. The proportional increase in carbohydrates with parasite virulence may therefore represent a behavioural adaptation by the host to fuel investment in immunity.

Finally, parasite treatment also influenced baseline lipid reserves **(Fig. 2c)**, with mosquitoes infected by selected parasites exhibiting higher lipid levels. The cause of this is unclear, but it may reflect greater parasite growth in these treatments compared to the unselected *Stock* parasite, leading to more lipid incorporation into spore production. It is also worth noting that our previous work showed that lipids are heavily exploited by the parasite later in infection [46], after day 22 of adulthood - beyond the time points measured here. Hence, lipid usage by the parasite may not yet have been strong enough to be detectable during the sampled stages, and due to the considerable mortality in some of the treatments, we could not measure later time points.

### (b) V. culicis infection alters host zinc and manganese levels independently of parasite selection

We observed notable changes in host metal content during infection **(Fig. 3)** and its usage by the parasite **(Fig. 4)**. Among the metals measured, only zinc **(Fig. 3b)** and manganese **(Fig. 3e)** were significantly affected by infection. Infections with any of the parasite treatments led to a decrease in host zinc content. This could reflect heavy utilization of zinc by the microsporidian parasite or represent a mechanism of immune evasion [44]. Zinc is essential for parasite growth and its proper metabolism [10,33], and is often linked to the production of virulence factors [65–67]. Furthermore, some fungal parasites can manipulate host immunity by inducing localised zinc deficits that impair immune responses [11,67–69]. However, our selection protocol had no detectable impact on zinc-related processes, as no differences were observed between parasite treatments.

In contrast, manganese levels increased following infection with any of the parasite lines **(Fig. 3e)**. This may indicate a parasite response to host immune defences, as many microsporidian antioxidant enzymes, such as manganese superoxide dismutases, depend on manganese for their activity and are particularly relevant for this taxon [70–72]. As with zinc, this increase in manganese was independent of the parasite’s selection history.

Across all treatments, we also noted a slight temporal increase in metal concentrations. Although the underlying mechanisms remain unclear, metals may become more bioavailable as infection progresses, potentially benefiting the host, the parasite, or both **(Fig. 3)**. From the parasite’s perspective, enhanced spore production over time would increase demand for metals sequestered from the host, making their accumulation more detectable at later stages. Future studies will be essential to better understand metal mobilization and dynamics during microsporidian infection and within the microsporidian cell.

### (c) Iron availability enhances parasite growth and spore production

Given the crucial role of metals in infection dynamics, and particularly the known importance of iron for parasite development, we next focused specifically on how iron availability affects *V. culicis* growth within the *A. gambiae* host.

Iron plays a vital role as a nutrient for growth in many parasite species [16,17,21,24,39,73], and it is particularly abundant in mosquitoes due to their hematophagous ecology [23,74]. In fungal parasites, iron acquisition is a critical determinant of virulence [75,76]. Many fungi have evolved sophisticated mechanisms to overcome host-imposed iron limitation, including the secretion of siderophores - small, high-affinity iron-transporter compounds - or the expression of more specialized iron transporter systems [18,77,78]. Although microsporidia lack many canonical metabolic pathways due to their highly reduced genomes [31], recent genomic studies suggest they have retained or adapted specific mechanisms for iron acquisition, including transporters likely involved in scavenging iron from the host cytoplasm [79], which was not the focus of the previously conducted metallomics.

Hence, in our study, we experimentally manipulated host iron levels by supplementing or chelating circulating iron in the mosquito haemolymph, consequently increasing and reducing its absorption by host cells and the replicating parasitic cells inside them. We then assessed the growth of different *V. culicis* treatments **(Fig. 4)**. Five-day iron supplementation and chelation led to an increase and decrease, respectively, in the production of infective spores at days 5 and 10 **(Fig. 4ab)**, but not at day 15, by which all mosquitoes harboured spores **(Fig. 4c)**. These effects were mirrored in parasite burden under the two dietary treatments **(Fig. 4d-f)**.

Overall, selected parasites exhibited a greater capacity to utilise circulating iron within the mosquito compared to the unselected *Stock*, resulting in accelerated production of infective spores. This result was not evident in the results from the metallomics, likely due to the small fraction of iron diverted to the parasite compared to the whole-body iron content that is not only circulating but also incorporated into many host cell structures. Differences in iron usage likely reflect adaptations arising during the selection experiment, favouring parasites that more efficiently sequester and use host iron. Enhanced iron acquisition is a known evolutionary strategy among intracellular parasites to sustain rapid proliferation under resource-limited conditions. However, the extent of these differences appeared inversely related to parasite growth: *Stock* parasites, initially with lower burdens, showed a larger relative increase in response to iron supplementation than selected parasites, which already maintained high burdens without supplementation. This trend was evident at day 15 **(Fig. 4f)**, when the parasite usually reaches its chronic persistent load, and suggests the lower increase in selected lines might be due to either limiting costs in the assimilation of iron by the parasite or limited evolvability at this time for the mechanism that is allowing a better iron sequestration by the parasite.

Interestingly, iron manipulation by fungal parasites is not only associated with growth but also with evasion of host immune responses [16,75,76]. In particular, the ability to modulate host iron homeostasis can impair the oxidative burst or other immune functions that are heavily dependent on iron-regulated pathways [80]. Whether microsporidia, including *V. culicis*, directly manipulate mosquito iron homeostasis or simply compete with the host for this resource remains an open question. Nevertheless, our findings provide evidence that iron availability is a critical driver of microsporidian growth within the host, and that virulence evolution in *V. culicis* may be partly linked to improved iron sequestration strategies.

### (d) Conclusion and perspectives

In conclusion, this study provides important insights into how *V. culicis* manipulates its host, *A. gambiae*, to exploit host resources for its growth and reproduction. We have shown that the parasite alters the dynamics of proteins, carbohydrates, lipids, and metals in a manner that is strongly influenced by the virulence of the parasite. More virulent strains of *V. culicis* seem to induce stronger immune responses in the host, resulting in higher protein levels and altered energy metabolism. Additionally, the study highlights the critical role of metal homeostasis, particularly zinc, manganese, and iron, in this host-parasite interaction, with iron availability being a key factor in parasite growth and spore production. These findings underscore the complex interplay between parasite, virulence, host resource allocation, and immune response.

Furthermore, the results suggest that the evolutionary adaptation of *V. culicis* to more efficiently exploit host resources, including iron, may play a significant role in driving within-host virulence and consequent host exploitation strategies. As the role of microsporidia as potential biological control agents in vector biology continues to be explored, these insights have implications for understanding the dynamics of parasitic disease and the potential for manipulating these interactions for disease control. Particularly concerning malaria, our findings suggest the inhibitory effect of microsporidian parasites on *Plasmodium spp.* might be due to competition for a common resource, rather than collateral immunity.

## Author contributions

LMS conceived the overall idea, JCK and LMS designed the experiments, LMS collected and analysed the data, AV performed the ICP-OES elemental analysis, and LMS wrote the first draft of the manuscript. All authors contributed critically to the drafts.

## Acknowledgements

We thank Tiago G. Zeferino and Gaëtan Glauser for their advice and technical support. LMS and the project were supported by SNF grant 310030_192786.

## Supplementary Information

**Table S1.**
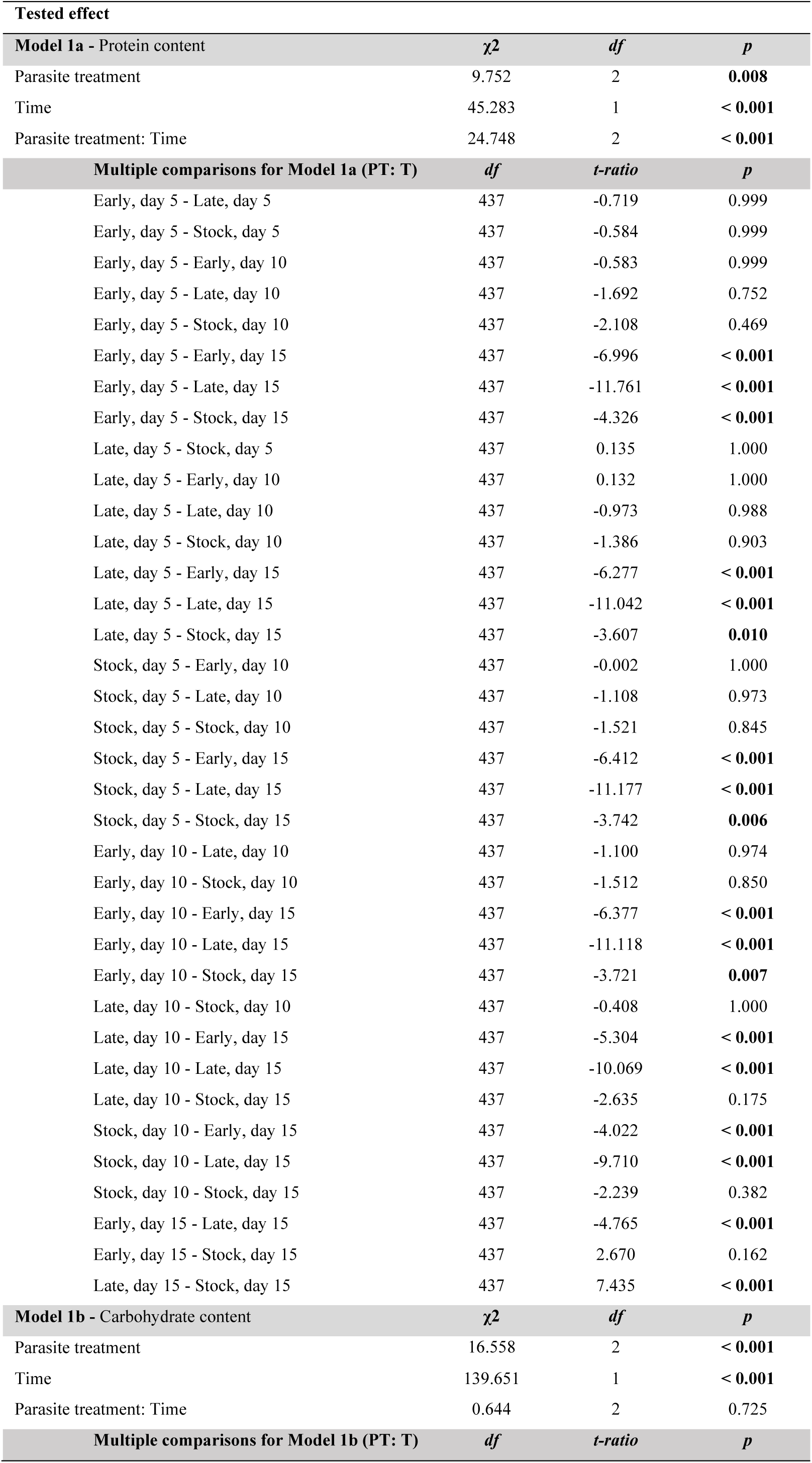
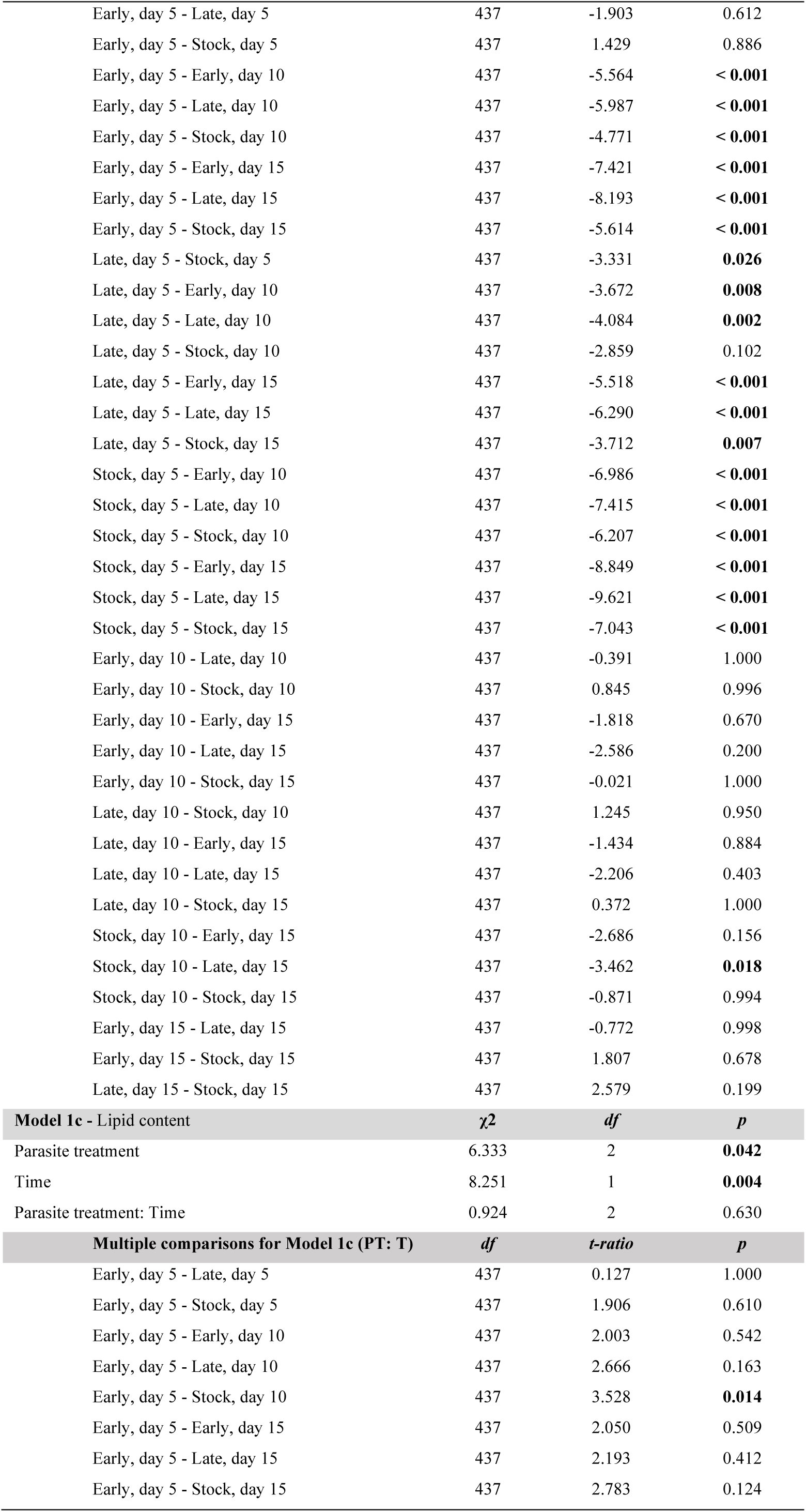
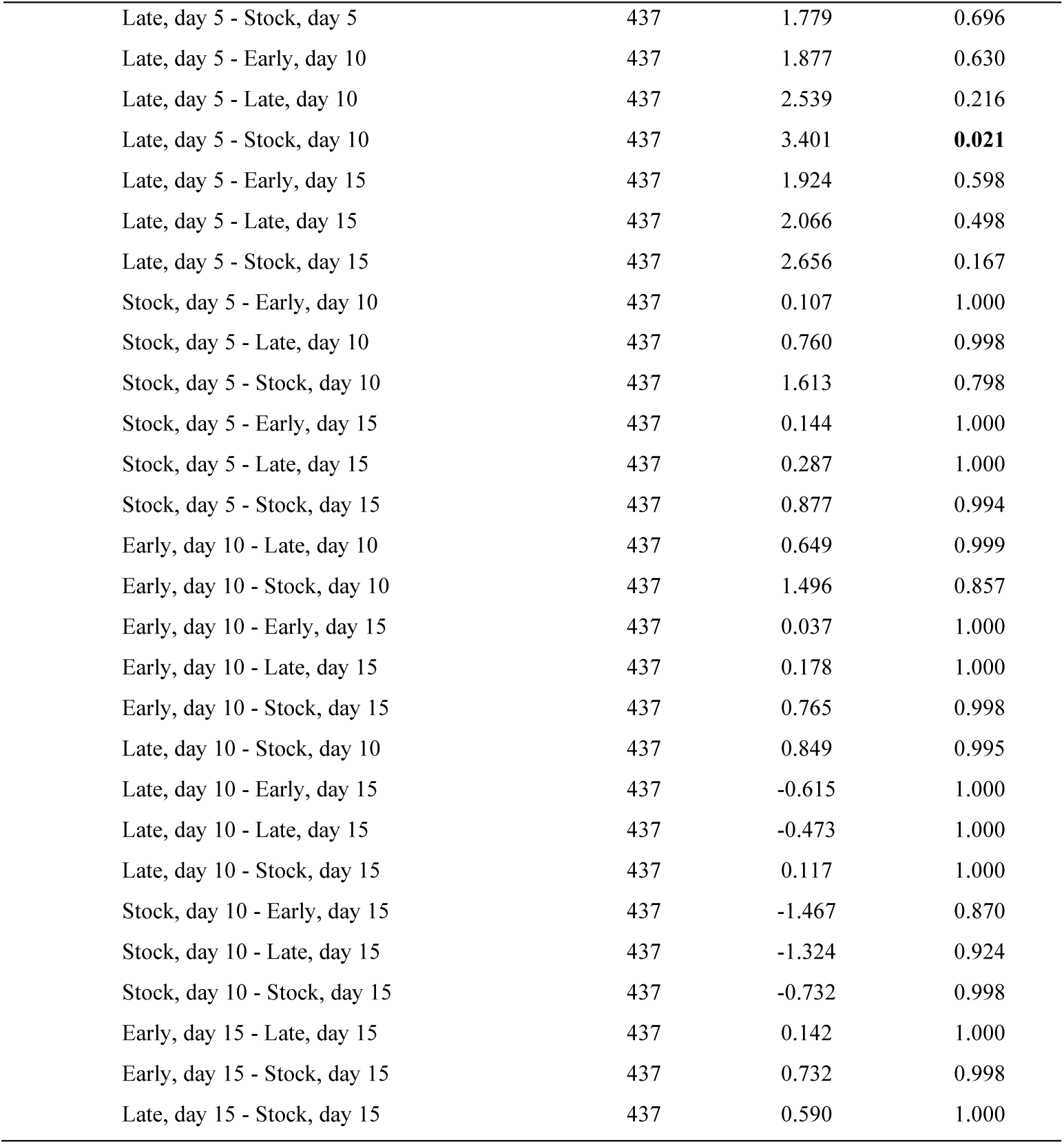
Protein, carbohydrate and lipid content through infection. Each resource was analysed independently. Differences in resource content were assessed using a linear mixed model for each of the three resources (proteins, carbohydrates, and lipids), with parasite treatment and time of collection as explanatory variables and their interaction. Replicate was considered a random factor. Individual resource content was quantified by measuring the sample’s quantity and adjusting it according to the respective individual weight. The square root of each resource content was calculated to fit the model best. Multiple comparisons tests were conducted to examine the interaction between “parasite treatment” and “time” and show pairwise differences between all combinatorial treatments.

**Table S2.**
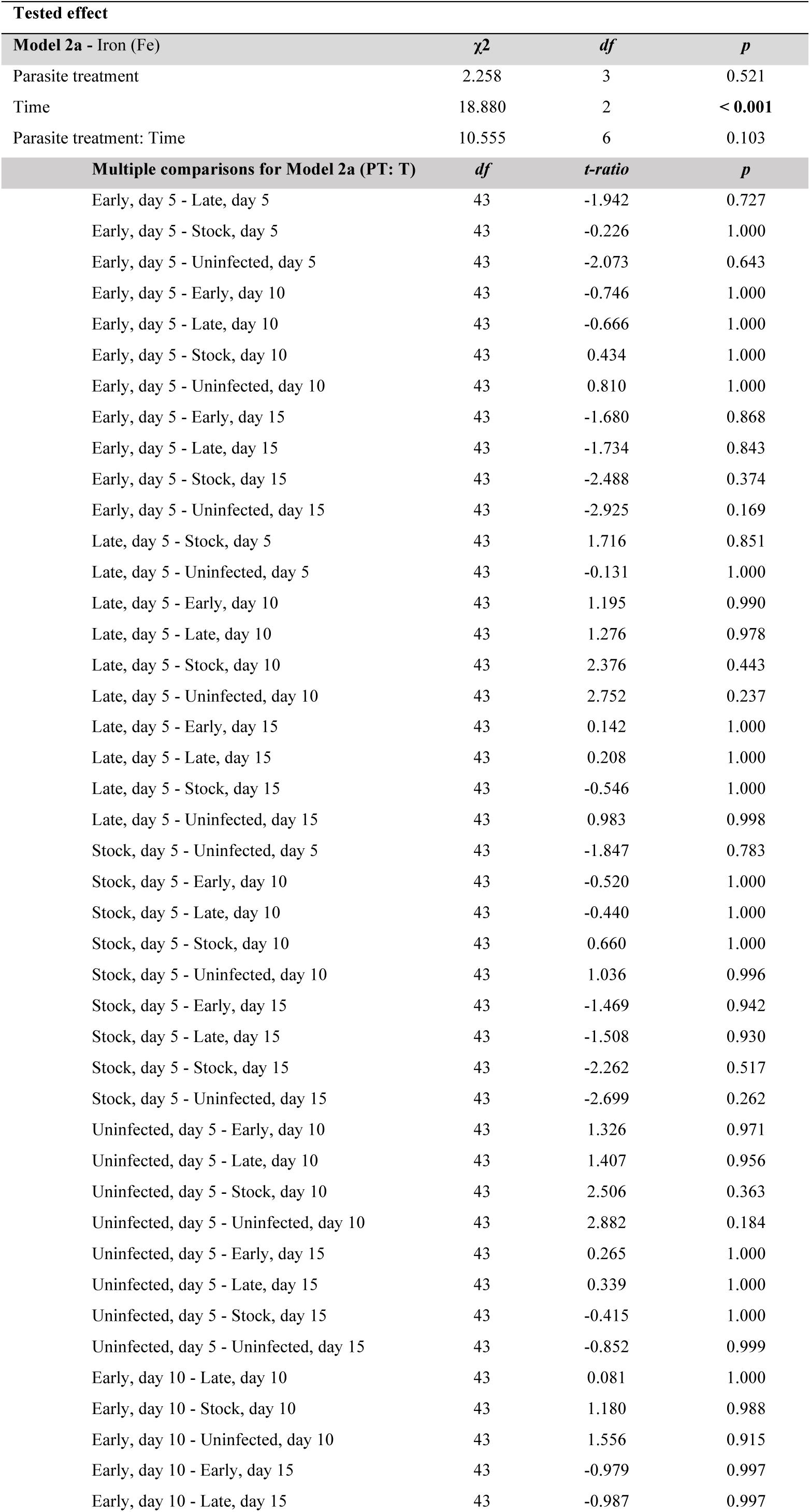
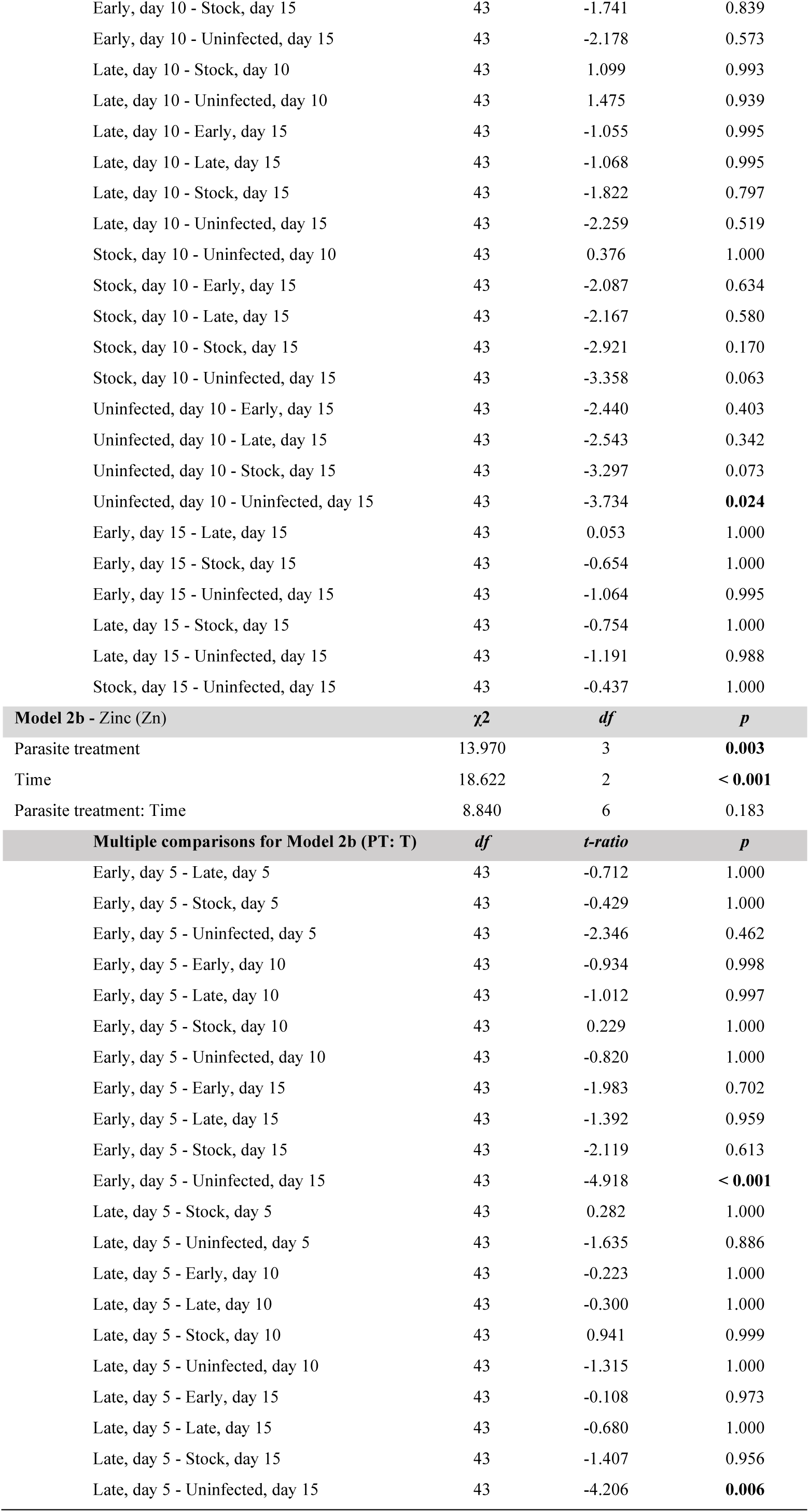
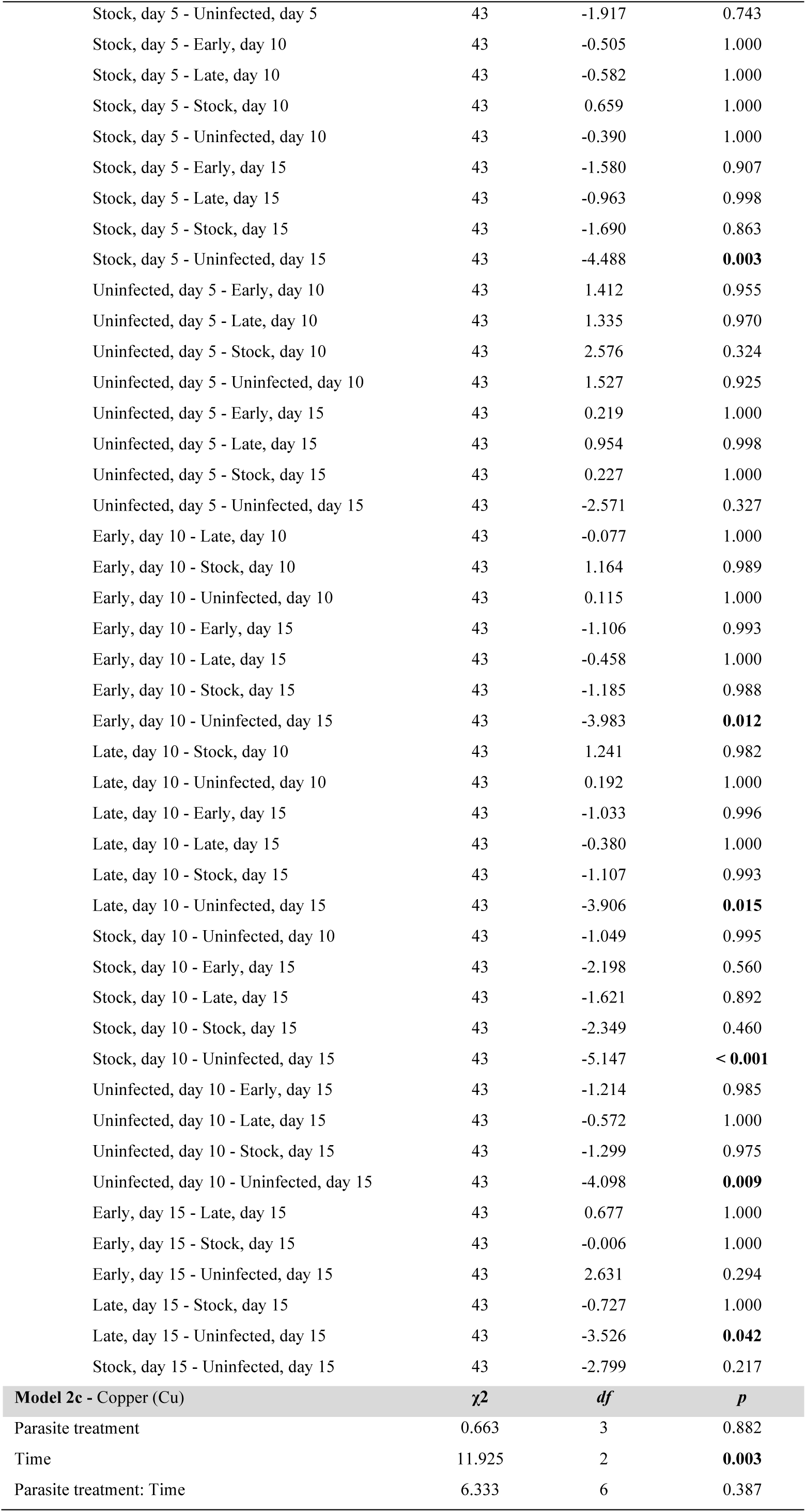
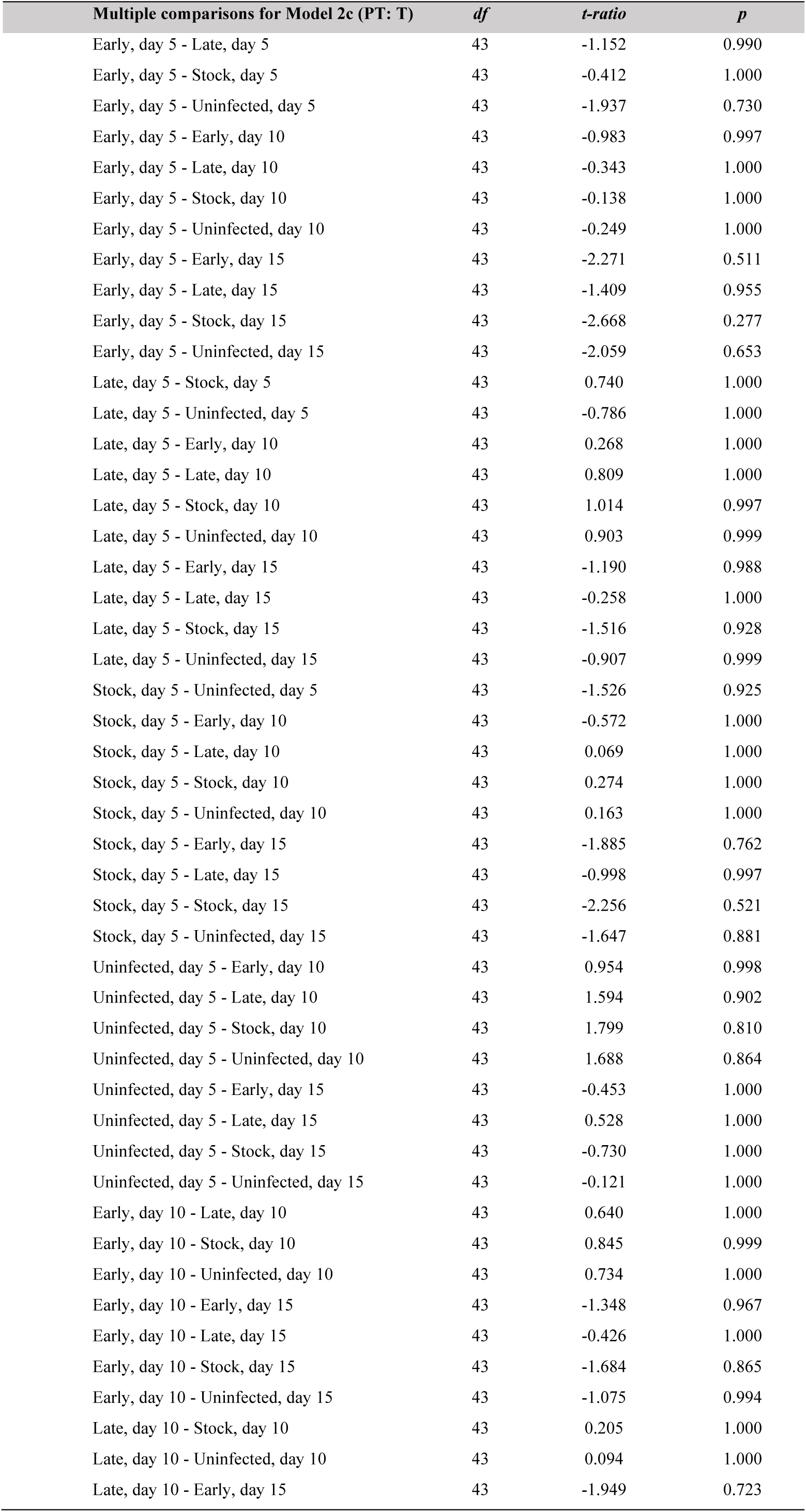
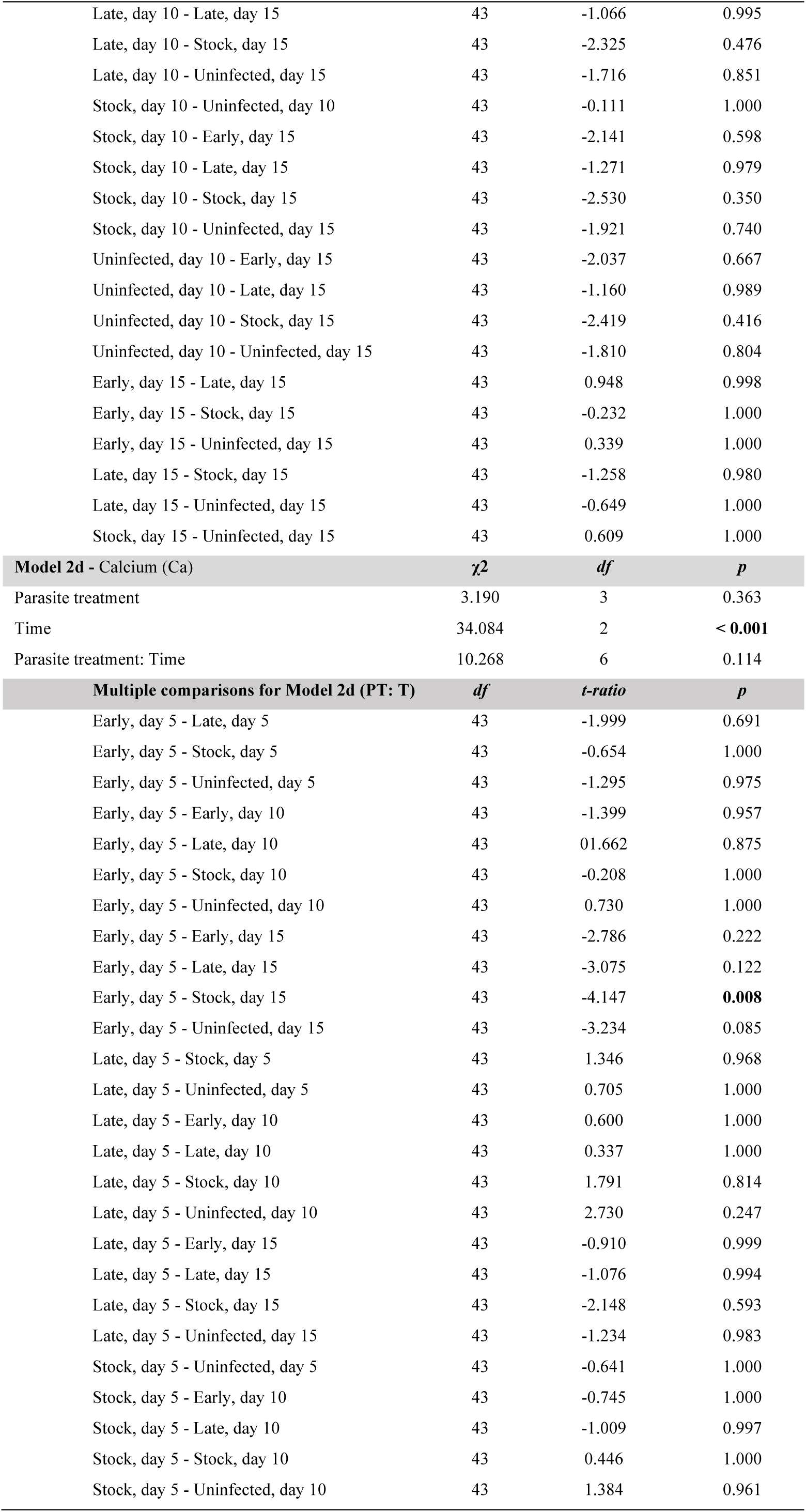
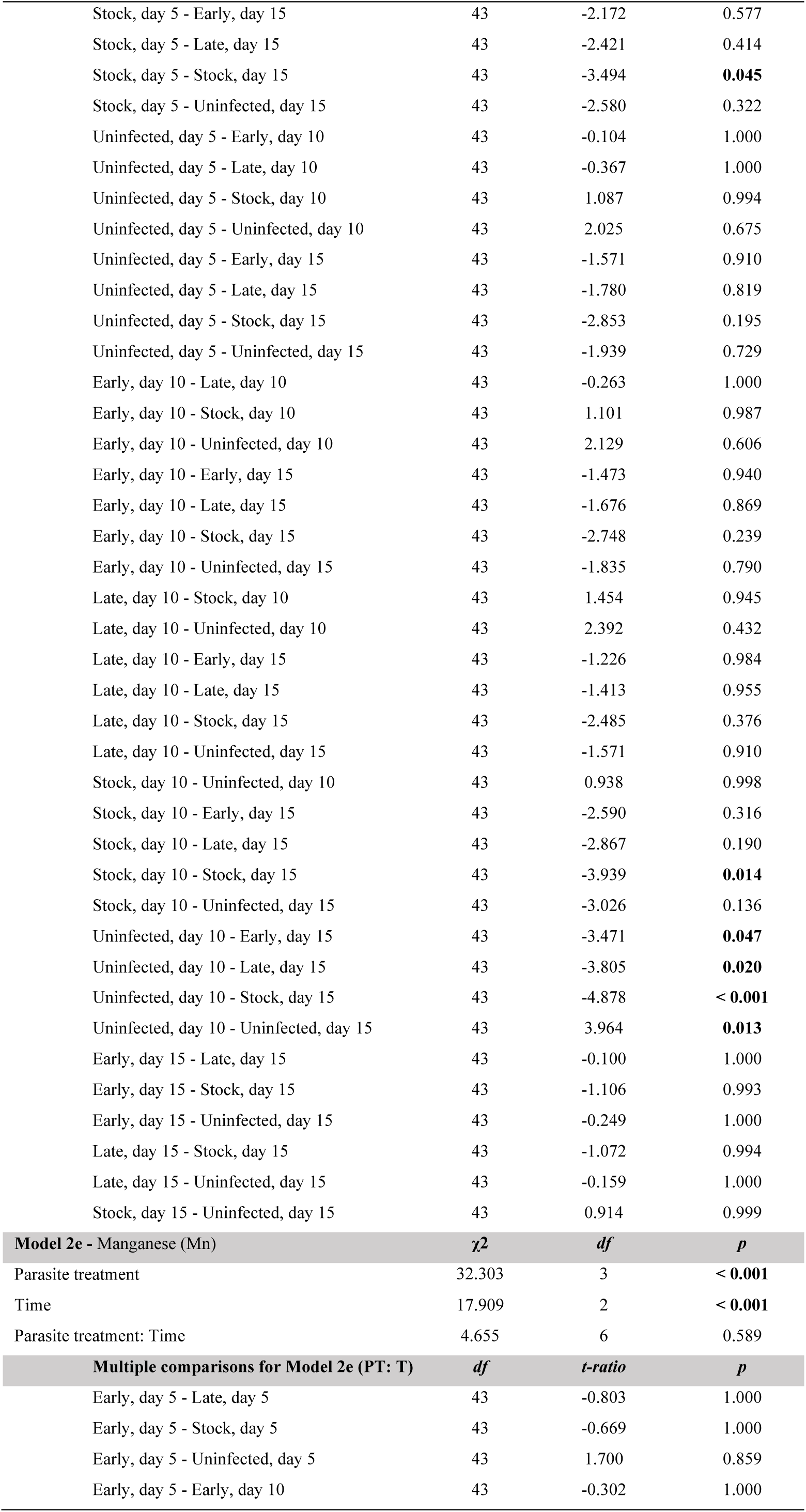
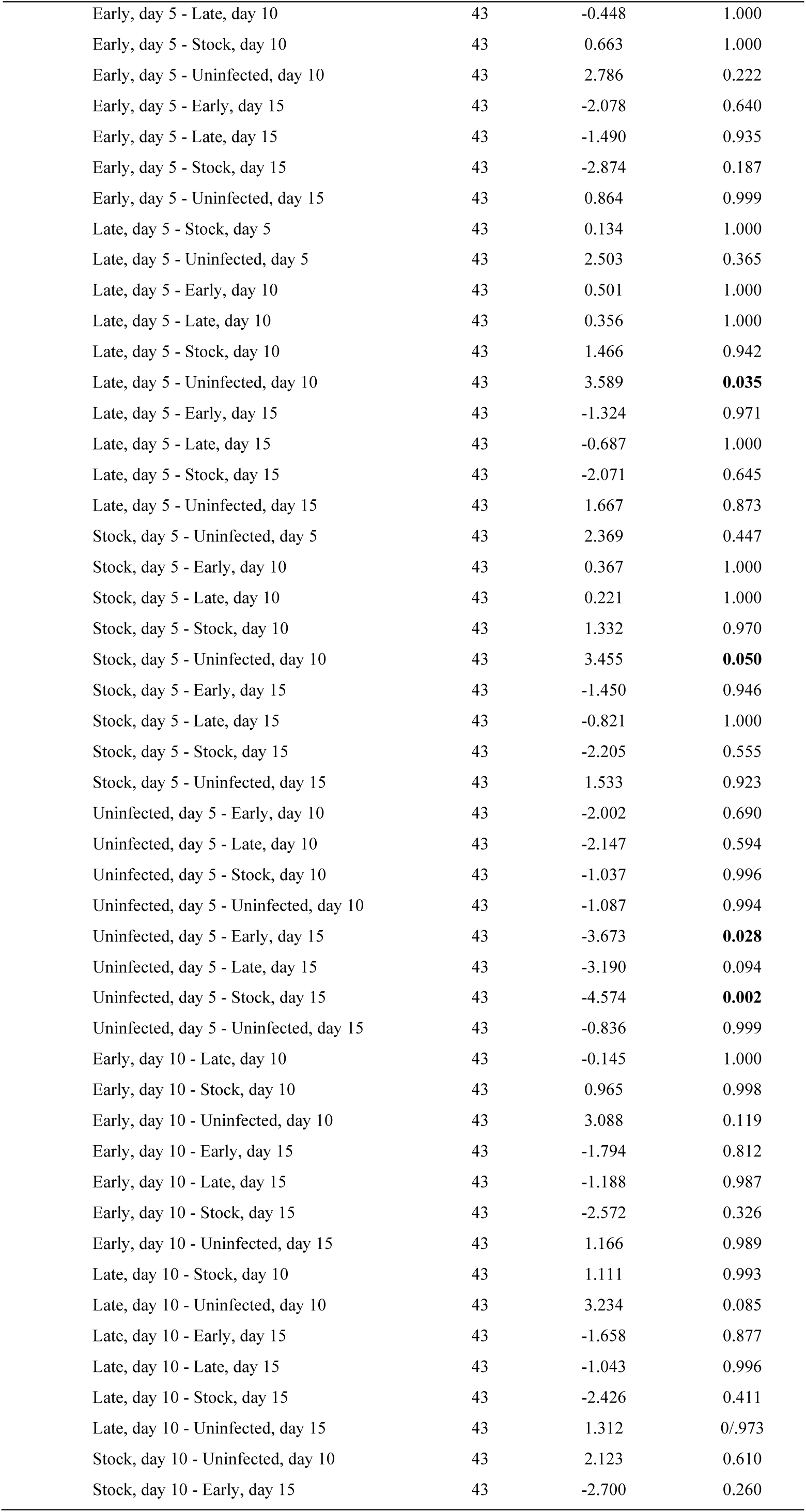
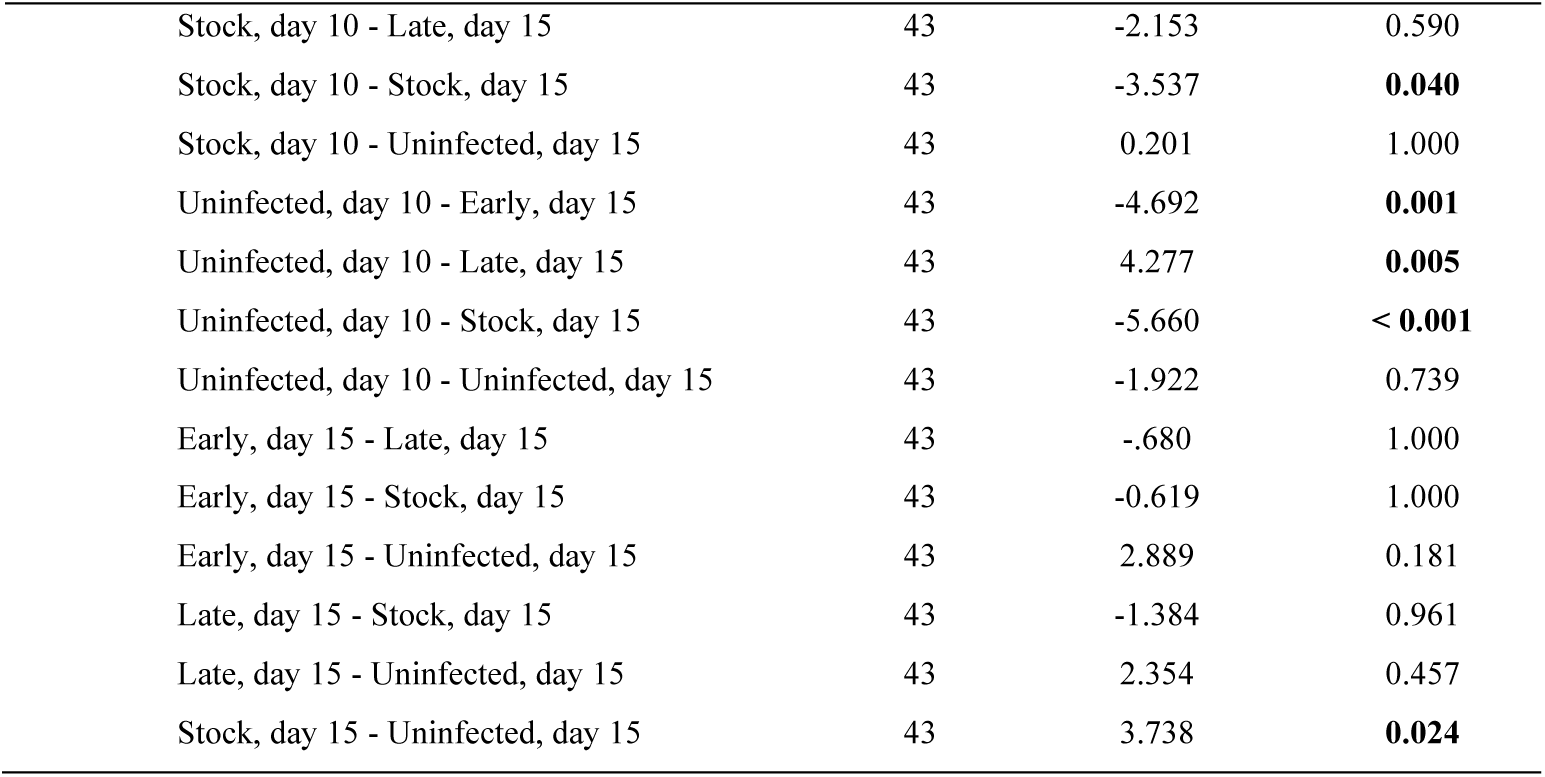
Metallomic profiles at adulthood. Similarly to the previous resources, each metal was analysed independently. We ran a linear mixed model for each metal element to test for differences in the metallomes of differently infected or uninfected mosquitoes across time. The dependent variable was the log-transformed concentration of the element obtained by ICP-OES, while the parasite treatment and assay day were explanatory variables and their interaction. Replicate was included as a random factor. Multiple comparisons tests were conducted to examine the interaction between “parasite treatment” and “time” and show pairwise differences between all combinatorial treatments.

**Table S3.** The effect of iron-supplementation and -chelation on parasite growth. We assessed the effect of the parasite treatment and diet on spore rate and growth for each time point assayed. For each spore rate comparison, we ran a generalised linear mixed model with a binomial error structure, where parasite treatment and diet treatment were set as explanatory variables, their interaction, and the presence or absence of spores as a response variable. Replicate was included as a random factor. Concerning spore growth analyses, we only considered individuals with at least one spore. We then ran a linear mixed model with the same explanatory and random factors as the previous one. Multiple comparisons tests were conducted to examine the interaction between “parasite treatment” and “diet” and show pairwise differences between all combinatorial treatments.

| Tested effect |  |  |  |
| --- | --- | --- | --- |
| Model 3a - Spore rate at day 5 | $\chi^2$ | df | p |
| Diet treatment | 15.430 | 2 | < 0.001 |
| Parasite treatment | 15.546 | 2 | < 0.001 |
| Diet treatment: Parasite treatment | 4.986 | 4 | 0.289 |
| Multiple comparisons for Model 3a (PT: D) | df | t-ratio | p |
| Early, Sugar - Late, Sugar | inf | -1.396 | 0.900 |
| Early, Sugar - Stock, Sugar | inf | 0.203 | 1.000 |
| Early, Sugar - Early, Sugar + BPS | inf | 1.460 | 0.874 |
| Early, Sugar - Late, Sugar + BPS | inf | 0.000 | 1.000 |
| Early, Sugar - Stock, Sugar + BPS | inf | 1.460 | 0.874 |
| Early, Sugar - Early, Sugar + FeCl<sub>3</sub> | inf | -1.790 | 0.689 |
| Early, Sugar - Late, Sugar + FeCl<sub>3</sub> | inf | -2.976 | 0.072 |
| Early, Sugar - Stock, Sugar + FeCl<sub>3</sub> | inf | 0.823 | 0.996 |
| Late, Sugar - Stock, Sugar | inf | 1.594 | 0.808 |
| Late, Sugar - Early, Sugar + BPS | inf | 2.792 | 0.118 |
| Late, Sugar - Late, Sugar + BPS | inf | 1.396 | 0.900 |
| Late, Sugar - Stock, Sugar + BPS | inf | 2.792 | 0.118 |
| Late, Sugar - Early, Sugar + FeCl<sub>3</sub> | inf | -0.405 | 1.000 |
| Late, Sugar - Late, Sugar + FeCl<sub>3</sub> | inf | -1.656 | 0.773 |
| Late, Sugar - Stock, Sugar + FeCl<sub>3</sub> | inf | 2.192 | 0.411 |
| Stock, Sugar - Early, Sugar + BPS | inf | 1.262 | 0.942 |
| Stock, Sugar - Late, Sugar + BPS | inf | -0.203 | 1.000 |
| Stock, Sugar - Stock, Sugar + BPS | inf | 1.262 | 0.942 |
| Stock, Sugar - Early, Sugar + FeCl<sub>3</sub> | inf | -1.986 | 0.553 |
| Stock, Sugar - Late, Sugar + FeCl<sub>3</sub> | inf | -3.160 | 0.042 |
| Stock, Sugar - Stock, Sugar + FeCl<sub>3</sub> | inf | 0.621 | 1.000 |
| Early, Sugar + BPS - Late, Sugar + BPS | inf | -1.460 | 0.874 |
| Early, Sugar + BPS - Stock, Sugar + BPS | inf | 0.000 | 1.000 |
| Early, Sugar + BPS - Early, Sugar + FeCl<sub>3</sub> | inf | -3.160 | 0.042 |
| Early, Sugar + BPS - Late, Sugar + FeCl<sub>3</sub> | inf | -4.241 | < 0.001 |
| Early, Sugar + BPS - Stock, Sugar + FeCl<sub>3</sub> | inf | -0.648 | 0.999 |
| Late, Sugar + BPS - Stock, Sugar + BPS | inf | 1.460 | 0.874 |
| Late, Sugar + BPS - Early, Sugar + FeCl<sub>3</sub> | inf | -1.790 | 0.689 |
| Late, Sugar + BPS - Late, Sugar + FeCl<sub>3</sub> | inf | -2.976 | 0.072 |
| Late, Sugar + BPS - Stock, Sugar + FeCl<sub>3</sub> | inf | 0.823 | 0.996 |
| Stock, Sugar + BPS - Early, Sugar + FeCl<sub>3</sub> | inf | -3.160 | 0.042 |
| Stock, Sugar + BPS - Late, Sugar + FeCl<sub>3</sub> | inf | -4.241 | < 0.001 |
| Stock, Sugar + BPS - Stock, Sugar + FeCl<sub>3</sub> | inf | -0.648 | 0.999 |
| Early, Sugar + FeCl<sub>3</sub> - Late, Sugar + FeCl<sub>3</sub> | inf | -1.262 | 0.942 |
| Early, Sugar + FeCl<sub>3</sub> - Stock, Sugar + FeCl<sub>3</sub> | inf | 2.574 | 0.200 |
| Late, Sugar + FeCl<sub>3</sub> - Stock, Sugar + FeCl<sub>3</sub> | inf | 3.705 | 0.007 |
| Model 3b- Spore rate at day 10 | $\chi^2$ | df | p |
| Diet treatment | 8.464 | 2 | **0.015** |
| Parasite treatment | 16.867 | 2 | **< 0.001** |
| Diet treatment: Parasite treatment | 0.504 | 4 | 0.973 |
| **Multiple comparisons for Model 3b (PT: D)** | **df** | **t-ratio** | **p** |
| Early, Sugar - Late, Sugar | inf | -0.001 | 1.000 |
| Early, Sugar - Stock, Sugar | inf | 1.275 | 0.9387 |
| Early, Sugar - Early, Sugar + BPS | inf | 2.209 | 0.3993 |
| Early, Sugar - Late, Sugar + BPS | inf | 0.972 | 0.9883 |
| Early, Sugar - Stock, Sugar + BPS | inf | 2.785 | 0.1195 |
| Early, Sugar - Early, Sugar + FeCl<sub>3</sub> | inf | -0.002 | 1.000 |
| Early, Sugar - Late, Sugar + FeCl<sub>3</sub> | inf | -0.002 | 1.000 |
| Early, Sugar - Stock, Sugar + FeCl<sub>3</sub> | inf | 0.575 | 1.000 |
| Late, Sugar - Stock, Sugar | inf | 0.001 | 1.000 |
| Late, Sugar - Early, Sugar + BPS | inf | 0.001 | 1.000 |
| Late, Sugar - Late, Sugar + BPS | inf | 0.001 | 1.000 |
| Late, Sugar - Stock, Sugar + BPS | inf | 0.001 | 1.000 |
| Late, Sugar - Early, Sugar + FeCl<sub>3</sub> | inf | 0.000 | 1.000 |
| Late, Sugar - Late, Sugar + FeCl<sub>3</sub> | inf | -0.002 | 1.000 |
| Late, Sugar - Stock, Sugar + FeCl<sub>3</sub> | inf | 0.001 | 0.878 |
| Stock, Sugar - Early, Sugar + BPS | inf | 1.451 | 1.000 |
| Stock, Sugar - Late, Sugar + BPS | inf | -0.391 | 0.252 |
| Stock, Sugar - Stock, Sugar + BPS | inf | 2.459 | 1.000 |
| Stock, Sugar - Early, Sugar + FeCl<sub>3</sub> | inf | -0.002 | 1.000 |
| Stock, Sugar - Late, Sugar + FeCl<sub>3</sub> | inf | -0.002 | 0.997 |
| Stock, Sugar - Stock, Sugar + FeCl<sub>3</sub> | inf | -0.826 | 0.706 |
| Early, Sugar + BPS - Late, Sugar + BPS | inf | -1.764 | 0.961 |
| Early, Sugar + BPS - Stock, Sugar + BPS | inf | 1.180 | 1.000 |
| Early, Sugar + BPS - Early, Sugar + FeCl<sub>3</sub> | inf | -0.002 | 1.000 |
| Early, Sugar + BPS - Late, Sugar + FeCl<sub>3</sub> | inf | -0.002 | 0.507 |
| Early, Sugar + BPS - Stock, Sugar + FeCl<sub>3</sub> | inf | -2.051 | 0.154 |
| Late, Sugar + BPS - Stock, Sugar + BPS | inf | 2.682 | 1.000 |
| Late, Sugar + BPS - Early, Sugar + FeCl<sub>3</sub> | inf | -0.002 | 1.000 |
| Late, Sugar + BPS - Late, Sugar + FeCl<sub>3</sub> | inf | -0.002 | 1.000 |
| Late, Sugar + BPS - Stock, Sugar + FeCl<sub>3</sub> | inf | -0.456 | 1.000 |
| Stock, Sugar + BPS - Early, Sugar + FeCl<sub>3</sub> | inf | -0.003 | 1.000 |
| Stock, Sugar + BPS - Late, Sugar + FeCl<sub>3</sub> | inf | -0.002 | 1.000 |
| Stock, Sugar + BPS - Stock, Sugar + FeCl<sub>3</sub> | inf | -2.837 | 0.105 |
| Early, Sugar + FeCl<sub>3</sub> - Late, Sugar + FeCl<sub>3</sub> | inf | -0.002 | 1.000 |
| Early, Sugar + FeCl<sub>3</sub> - Stock, Sugar + FeCl<sub>3</sub> | inf | 0.002 | 1.000 |
| Late, Sugar + FeCl<sub>3</sub> - Stock, Sugar + FeCl<sub>3</sub> | inf | 0.002 | 1.000 |
| **Model 3d - Spore load at day 5** | **χ<sup>2</sup>** | **df** | **p** |
| Diet treatment | 37.751 | 2 | **< 0.001** |
| Parasite treatment | 12.451 | 2 | **0.002** |
| Diet treatment: Parasite treatment | 6.925 | 4 | 0.140 |
| **Multiple comparisons for Model 3d (PT: D)** | **df** | **t-ratio** | **p** |
| Early, Sugar - Late, Sugar | 190 | -1.257 | 0.942 |
| Early, Sugar - Stock, Sugar | 189 | 1.284 | 0.935 |
| Early, Sugar - Early, Sugar + BPS | 190 | 1.844 | 0.653 |
| Early, Sugar - Late, Sugar + BPS | 190 | 0.494 | 1.000 |
| Early, Sugar - Stock, Sugar + BPS | 190 | 2.225 | 0.394 |
| Early, Sugar - Early, Sugar + FeCl<sub>3</sub> | 188 | 0.558 | 1.000 |
| Early, Sugar - Late, Sugar + FeCl<sub>3</sub> | 189 | -3.197 | **0.042** |
| Early, Sugar - Stock, Sugar + FeCl<sub>3</sub> | 189 | 2.128 | 0.458 |
| Late, Sugar - Stock, Sugar | 189 | 2.613 | 0.188 |
| Late, Sugar - Early, Sugar + BPS | 192 | 3.041 | 0.066 |
| Late, Sugar - Late, Sugar + BPS | 188 | 1.791 | 0.689 |
| Late, Sugar - Stock, Sugar + BPS | 190 | 3.456 | **0.019** |
| Late, Sugar - Early, Sugar + FeCl<sub>3</sub> | 190 | 1.982 | 0.558 |
| Late, Sugar - Late, Sugar + FeCl<sub>3</sub> | 189 | -2.040 | 0.518 |
| Late, Sugar - Stock, Sugar + FeCl<sub>3</sub> | 189 | 3.439 | **0.020** |
| Stock, Sugar - Early, Sugar + BPS | 190 | 0.676 | 0.999 |
| Stock, Sugar - Late, Sugar + BPS | 189 | -0.795 | 0.997 |
| Stock, Sugar - Stock, Sugar + BPS | 190 | 1.057 | 0.980 |
| Stock, Sugar - Early, Sugar + FeCl<sub>3</sub> | 188 | -0.841 | 0.995 |
| Stock, Sugar - Late, Sugar + FeCl<sub>3</sub> | 188 | -4.591 | **< 0.001** |
| Stock, Sugar - Stock, Sugar + FeCl<sub>3</sub> | 188 | 0.889 | 0.993 |
| Early, Sugar + BPS - Late, Sugar + BPS | 191 | -1.398 | 0.898 |
| Early, Sugar + BPS - Stock, Sugar + BPS | 189 | 0.351 | 1.000 |
| Early, Sugar + BPS - Early, Sugar + FeCl<sub>3</sub> | 190 | -1.476 | 0.865 |
| Early, Sugar + BPS - Late, Sugar + FeCl<sub>3</sub> | 190 | -4.790 | **< 0.001** |
| Early, Sugar + BPS - Stock, Sugar + FeCl<sub>3</sub> | 190 | 0.157 | 1.000 |
| Late, Sugar + BPS - Stock, Sugar + BPS | 189 | 1.786 | 0.691 |
| Late, Sugar + BPS - Early, Sugar + FeCl<sub>3</sub> | 190 | 0.020 | 1.000 |
| Late, Sugar + BPS - Late, Sugar + FeCl<sub>3</sub> | 190 | -3.753 | **0.007** |
| Late, Sugar + BPS - Stock, Sugar + FeCl<sub>3</sub> | 189 | 1.660 | 0.770 |
| Stock, Sugar + BPS - Early, Sugar + FeCl<sub>3</sub> | 190 | -1.883 | 0.626 |
| Stock, Sugar + BPS - Late, Sugar + FeCl<sub>3</sub> | 190 | -5.217 | **< 0.001** |
| Stock, Sugar + BPS - Stock, Sugar + FeCl<sub>3</sub> | 190 | -0.210 | 1.000 |
| Early, Sugar + FeCl<sub>3</sub> - Late, Sugar + FeCl<sub>3</sub> | 189 | -4.193 | **0.001** |
| Early, Sugar + FeCl<sub>3</sub> - Stock, Sugar + FeCl<sub>3</sub> | 189 | 1.764 | 0.706 |
| Late, Sugar + FeCl<sub>3</sub> - Stock, Sugar + FeCl<sub>3</sub> | 189 | 5.343 | **< 0.001** |
| **Model 3e - Spore load at day 10** | **$\chi^2$** | ***df*** | ***p*** |
| Diet treatment | 37.601 | 2 | **< 0.001** |
| Parasite treatment | 48.705 | 2 | **< 0.001** |
| Diet treatment: Parasite treatment | 3.038 | 4 | 0.552 |
| **Multiple comparisons for Model 3e (PT: D)** | ***df*** | ***t-ratio*** | ***p*** |
| Early, Sugar - Late, Sugar | 404 | -3.455 | **0.018** |
| Early, Sugar - Stock, Sugar | 404 | 0.192 | 1.000 |
| Early, Sugar - Early, Sugar + BPS | 404 | 2.299 | 0.345 |
| Early, Sugar - Late, Sugar + BPS | 404 | -0.281 | 1.000 |
| Early, Sugar - Stock, Sugar + BPS | 404 | 1.247 | 0.945 |
| Early, Sugar - Early, Sugar + FeCl<sub>3</sub> | 404 | -2.241 | 0.381 |
| Early, Sugar - Late, Sugar + FeCl<sub>3</sub> | 404 | -5.341 | **< 0.001** |
| Early, Sugar - Stock, Sugar + FeCl<sub>3</sub> | 404 | -1.372 | 0.908 |
| Late, Sugar - Stock, Sugar | 404 | 3.592 | **0.048** |
| Late, Sugar - Early, Sugar + BPS | 404 | 5.606 | **< 0.001** |
| Late, Sugar - Late, Sugar + BPS | 404 | 3.137 | **0.048** |
| Late, Sugar - Stock, Sugar + BPS | 404 | 4.429 | **< 0.001** |
| Late, Sugar - Early, Sugar + FeCl<sub>3</sub> | 404 | 1.220 | 0.952 |
| Late, Sugar - Late, Sugar + FeCl<sub>3</sub> | 404 | -1.896 | 0.618 |
| Late, Sugar - Stock, Sugar + FeCl<sub>3</sub> | 404 | 2.058 | 0.504 |
| Stock, Sugar - Early, Sugar + BPS | 404 | 2.083 | 0.487 |
| Stock, Sugar - Late, Sugar + BPS | 404 | -0.466 | 1.000 |
| Stock, Sugar - Stock, Sugar + BPS | 404 | 1.054 | 0.980 |
| Stock, Sugar - Early, Sugar + FeCl<sub>3</sub> | 404 | -2.398 | 0.288 |
| Stock, Sugar - Late, Sugar + FeCl<sub>3</sub> | 404 | -5.448 | **< 0.001** |
| Stock, Sugar - Stock, Sugar + FeCl<sub>3</sub> | 404 | -1.541 | 0.835 |
| Early, Sugar + BPS - Late, Sugar + BPS | 404 | -2.546 | 0.214 |
| Early, Sugar + BPS - Stock, Sugar + BPS | 404 | -0.932 | 0.991 |
| Early, Sugar + BPS - Early, Sugar + FeCl<sub>3</sub> | 404 | -4.448 | **< 0.001** |
| Early, Sugar + BPS - Late, Sugar + FeCl<sub>3</sub> | 404 | -7.405 | **< 0.001** |
| Early, Sugar + BPS - Stock, Sugar + FeCl<sub>3</sub> | 404 | -3.599 | **0.011** |
| Late, Sugar + BPS - Stock, Sugar + BPS | 404 | 1.495 | 0.858 |
| Late, Sugar + BPS - Early, Sugar + FeCl<sub>3</sub> | 404 | -1.935 | 0.589 |
| Late, Sugar + BPS - Late, Sugar + FeCl<sub>3</sub> | 404 | -5.002 | **< 0.001** |
| Late, Sugar + BPS - Stock, Sugar + FeCl<sub>3</sub> | 404 | -1.078 | 0.977 |
| Stock, Sugar + BPS - Early, Sugar + FeCl<sub>3</sub> | 404 | -3.313 | **0.028** |
| Stock, Sugar + BPS - Late, Sugar + FeCl<sub>3</sub> | 404 | -6.163 | **< 0.001** |
| Stock, Sugar + BPS - Stock, Sugar + FeCl<sub>3</sub> | 404 | -2.5-6 | 0.232 |
| Early, Sugar + FeCl<sub>3</sub> - Late, Sugar + FeCl<sub>3</sub> | 404 | -3.116 | 0.050 |
| Early, Sugar + FeCl<sub>3</sub> - Stock, Sugar + FeCl<sub>3</sub> | 404 | -0.850 | 0.996 |
| Late, Sugar + FeCl<sub>3</sub> - Stock, Sugar + FeCl<sub>3</sub> | 404 | 3.934 | **0.003** |
| **Model 3f - Spore load at day 15** | **$\chi^2$** | ***df*** | ***p*** |
| Diet treatment | 7.271 | 2 | **0.0264** |
| Parasite treatment | 79.910 | 2 | **< 0.001** |
| Diet treatment: Parasite treatment | 18.102 | 4 | **0.001** |
| **Multiple comparisons for Model 3f (PT: D)** | ***df*** | ***t-ratio*** | ***p*** |
| Early, Sugar - Late, Sugar | 437 | -1.128 | 0.970 |
| Early, Sugar - Stock, Sugar | 437 | 1.553 | 0.830 |
| Early, Sugar - Early, Sugar + BPS | 437 | 7.691 | **< 0.001** |
| Early, Sugar - Late, Sugar + BPS | 437 | 2.497 | 0.236 |
| Early, Sugar - Stock, Sugar + BPS | 437 | 5.713 | **< 0.001** |
| Early, Sugar - Early, Sugar + FeCl<sub>3</sub> | 437 | -0.074 | 1.000 |
| Early, Sugar - Late, Sugar + FeCl<sub>3</sub> | 437 | -2.633 | 0.176 |
| Early, Sugar - Stock, Sugar + FeCl<sub>3</sub> | 437 | -3.095 | 0.053 |
| Late, Sugar - Stock, Sugar | 437 | 2.681 | 0.158 |
| Late, Sugar - Early, Sugar + BPS | 437 | 8.8219 | **< 0.001** |
| Late, Sugar - Late, Sugar + BPS | 437 | 3.625 | **0.010** |
| Late, Sugar - Stock, Sugar + BPS | 437 | 6.841 | **< 0.001** |
| Late, Sugar - Early, Sugar + FeCl<sub>3</sub> | 437 | 1.053 | 0.980 |
| Late, Sugar - Late, Sugar + FeCl<sub>3</sub> | 437 | -1.505 | 0.853 |
| Late, Sugar - Stock, Sugar + FeCl<sub>3</sub> | 437 | -1.967 | 0.567 |
| Stock, Sugar - Early, Sugar + BPS | 437 | 6.139 | **< 0.001** |
| Stock, Sugar - Late, Sugar + BPS | 437 | 0.944 | 0.990 |
| Stock, Sugar - Stock, Sugar + BPS | 437 | 4.160 | **0.001** |
| Stock, Sugar - Early, Sugar + FeCl<sub>3</sub> | 437 | -1.627 | 0.790 |
| Stock, Sugar - Late, Sugar + FeCl<sub>3</sub> | 437 | -4.186 | **0.001** |
| Stock, Sugar - Stock, Sugar + FeCl<sub>3</sub> | 437 | -4.648 | < 0.001 |
| Early, Sugar + BPS - Late, Sugar + BPS | 437 | -5.194 | < 0.001 |
| Early, Sugar + BPS - Stock, Sugar + BPS | 437 | -1.979 | 0.559 |
| Early, Sugar + BPS - Early, Sugar + FeCl<sub>3</sub> | 437 | -7.766 | < 0.001 |
| Early, Sugar + BPS - Late, Sugar + FeCl<sub>3</sub> | 437 | -10.325 | < 0.001 |
| Early, Sugar + BPS - Stock, Sugar + FeCl<sub>3</sub> | 437 | -10.787 | < 0.001 |
| Late, Sugar + BPS - Stock, Sugar + BPS | 437 | 3.216 | 0.037 |
| Late, Sugar + BPS - Early, Sugar + FeCl<sub>3</sub> | 437 | -2.572 | 0.202 |
| Late, Sugar + BPS - Late, Sugar + FeCl<sub>3</sub> | 437 | -5.130 | < 0.001 |
| Late, Sugar + BPS - Stock, Sugar + FeCl<sub>3</sub> | 437 | -5.592 | < 0.001 |
| Stock, Sugar + BPS - Early, Sugar + FeCl<sub>3</sub> | 437 | -5.787 | < 0.001 |
| Stock, Sugar + BPS - Late, Sugar + FeCl<sub>3</sub> | 437 | -8.346 | < 0.001 |
| Stock, Sugar + BPS - Stock, Sugar + FeCl<sub>3</sub> | 437 | -8.808 | < 0.001 |
| Early, Sugar + FeCl<sub>3</sub> - Late, Sugar + FeCl<sub>3</sub> | 437 | -2.559 | 0.207 |
| Early, Sugar + FeCl<sub>3</sub> - Stock, Sugar + FeCl<sub>3</sub> | 437 | -3.021 | 0.066 |
| Late, Sugar + FeCl<sub>3</sub> - Stock, Sugar + FeCl<sub>3</sub> | 437 | -0.462 | 1.000 |

## Notes

### Competing Interest Statement

The authors have declared no competing interest.

### Summary of Updates

New revisions from the peer reviewing process have been incorporated.

